# *singlecellVR*: interactive visualization of single-cell data in virtual reality

**DOI:** 10.1101/2020.07.30.229534

**Authors:** David F. Stein, Huidong Chen, Michael E. Vinyard, Qian Qin, Rebecca D. Combs, Qian Zhang, Luca Pinello

## Abstract

Single-cell assays have transformed our ability to model heterogeneity within cell populations. As these assays have advanced in their ability to measure various aspects of molecular processes in cells, computational methods to analyze and meaningfully visualize such data have required matched innovation. Independently, Virtual Reality (VR) has recently emerged as a powerful technology to dynamically explore complex data and shows promise for adaptation to challenges in single-cell data visualization. However, adopting VR for single-cell data visualization has thus far been hindered by expensive prerequisite hardware or advanced data preprocessing skills. To address current shortcomings, we present *singlecellVR*, a user-friendly web application for visualizing single-cell data, designed for cheap and easily available virtual reality hardware (e.g., Google Cardboard, ∼$10). *singlecellVR* can visualize data from a variety of sequencing-based technologies including transcriptomic, epigenomic, and proteomic data as well as combinations thereof. Analysis modalities supported include approaches to clustering as well as trajectory inference and visualization of dynamical changes discovered through modelling RNA velocity. We provide a companion software package, *scvr* to streamline data conversion from the most widely-adopted single-cell analysis tools as well as a growing database of pre-analyzed datasets to which users can contribute.

## 1 Introduction

Characterization of cell type, while once dominated by pathological description, has over the past decade shifted towards a more quantitative and molecular approach. As such, molecular measurements in single cells have emerged as the centerpiece of the current paradigm of mechanistic biological investigation (Trapnell, 2015). Technological advancements have enabled researchers to measure all aspects of the central dogma of molecular biology at the single-cell level (Stuart and Satija, 2019). Single-cell RNA sequencing (scRNA-seq), a technique that profiles the relative expression of genes in individual cells and single-cell Assay for Transposase Accessible Chromatin using sequencing (scATAC-seq), a technique that surveys genome-wide chromatin accessibility are the most well-established and widely-used of these methods (Buenrostro et al., 2015; Lähnemann et al., 2020). In fact, combined scRNA-seq + scATAC-seq assays are now routine (Perkel, 2021). Additionally, assays to profile DNA methylation (Luo et al., 2018) or protein levels are now maturing and becoming more widely-accessible (Specht et al., 2019; Labib and Kelley, 2020). Most recently, combinations of various data modalities can now routinely be collected in parallel from the same cell(Chen et al., 2019d; Zhu et al., 2019; Ma et al., 2020; Xing et al., 2020; Swanson et al., 2021).

scRNA-seq experiments generate on the order of millions of sequencing reads that sample the relative expression of approximately 20,000 – 30,000 transcribed features (e.g., genes) in each cell of the sample. Normalized read counts for each feature can then be compared to discern differences between cells. scATAC-seq samples comprise a larger feature space wherein cells are characterized by the genomic coordinates of chromatin accessible regions and sequence-features derived from these regions (e.g. transcription factor motifs, *k*-mer frequencies, etc.). Initially performed in dozens to hundreds of cells, these experiments are now performed on the order of millions of cells. With a high dimensional feature space as a result of thousands of features being considered for each cell and large (in cell number) experiments, analysis methods for this data have been required to advance concurrently with the development of these technologies (Chen et al., 2019c; Tian et al., 2019).

With the exception of proof-of-concept methods still too nascent to be widely applied (Chen et al., 2021), omics measurements of single-cells are generally destructive, preventing measurement of a cell at more than a single timepoint. As a result, most single-cell measurements for studying dynamic processes is of a “snapshot” nature, imposing inherent limitations on the study of such processes from this data (Weinreb et al., 2018). In light of this, transcription rates can be informative of ongoing processes in cells. The recent advent of *RNA velocity* quantifies and models the ratios of spliced and un-spliced RNA (mRNA and pre-mRNA, respectively) such that they indicate the temporal derivative of gene expression patterns and thereby reflect dynamic cellular processes, allowing predictions of past and future cell states(La Manno et al., 2018).

Among others, PCA, t-SNE, and UMAP are dimensional reduction methods that have become common choices for enabling the visualization of high-dimensional single-cell datasets. Dimensionally reduced datasets are plotted such that similar cells cluster together and those with highly differing features are likewise clustered apart. In addition to the visualization and clustering of cells, trajectory inference methods have been proposed to learn a latent topological structure to reconstruct the putative time-ordering (*pseudotime*) by which cells may progress along a dynamic biological process (Saelens et al., 2018). As single-cell technologies have advanced, techniques to cluster and organize cells based on single-cell assays have advanced alongside them, allowing key insights toward cell type and state characterization. Combined with RNA velocity information, trajectory inference can offer key insights on dynamical changes to cell states. Once in press however, representation of these dimensionally-reduced visualizations is limited to just two or three dimensions. Even using three-dimensional plots from published studies, one cannot dynamically adjust or rotate the visualization to better understand the data from another angle. In addition, cells are typically annotated by features (e.g. time points, cell type or clusters) to investigate stratification along an axis of some biological process. To change the annotations presented in publication, one must often reprocess the raw data, which is time-and skill-intensive, highlighting the need for more dynamical visualization tools. While such current data representations are often limited and static, single-cell omic datasets are information-rich and, in many cases, important biological heterogeneity cannot be easily investigated or visualized outside the scope of the original publication, without spending considerable cost and time to reanalyze the datasets from scratch.

VR visualization methods for single cell data have been recently proposed (Yang et al., 2018; Legetth et al., 2019; Bressan et al., 2021). However, these methods require either expensive hardware or specific data inputs that mandate intermediate to advanced computational skills. Thus, tools and clear protocols are required to enable researchers, especially those who are not able to efficiently reprocess the raw data, to explore the richness of published datasets (or their own unpublished data) through a simple, easy and affordable VR platform. Importantly, this platform must be flexible enough to accept all types of omics data from established and emerging technologies and processing tools currently employed by the single cell community.

At the time of this writing, three non-peer-reviewed methods employing VR technology that produce two- and three-dimensional visualizations of single-cell data have recently been reported. *CellexalVR* enables the visualization of standard scRNA-seq data though requires users to preprocess their data through scripting (Legetth et al., 2019). Unfortunately, this tool also requires expensive and dedicated VR hardware to operate. Another recent method for visualizing single-cell data in VR is *Theia* (Bressan et al., 2021), which has been designed with a focus on the exploration of spatial datasets for both RNA and protein measurements. Similar to *CellexalVR*, expensive computing power and VR hardware required to use *Theia* creates a barrier to entry. An alternative to these high-performance methods for VR visualization of single-cell data is *starmap* (Yang et al., 2018), which allows the use of inexpensive cardboard visor hardware. However, *starmap* lacks the advanced portability of outputs from commonly-used scRNA-seq analysis tools and limits cell annotation to clustering results of transcriptomic data. Of note, there are currently no peer-reviewed tools available for the visualization of single-cell data in VR illustrating the novelty in this area of research. To overcome the limitations of these existing methods as well as build on their qualities and initial progress, we present *singlecellVR*, an interactive web application, which implements a flexible, innovative visualization for various modalities of single-cell data built on VR technology. *singlecellVR* supports clustering, trajectory inference and abstract graph analysis for transcriptomic as well as epigenomic and proteomic single cell data. Importantly, *singlecellVR* supports visualization of cell dynamics as described by RNA velocity, a recent milestone in the sequence-based analysis of single cells (La Manno et al., 2018; Bergen et al., 2020). *singlecellVR* is a browser-contained, free, and open-access tool. Notably, we have developed a one-command conversion tool, *scvr* to directly prepare the results of commonly-used single-cell analysis tools for visualization using *singlecellVR*.

## 2 Results

### 2.1 singlecellVR user experience and overview

*singlecellVR* is an easy-to-use web platform and database that can be operated from inexpensive, cardboard visor hardware that costs as little as ∼$10 and is available online from popular vendors including Google and Amazon. The webpage, available at https://www.singlecellvr.com enables users to explore several pre-loaded datasets or upload their own datasets for VR visualization. Visualization can be done either on a personal computer or smartphone. To facilitate the transition between the personal computer browser view and the phone-enabled VR visor (VR mode), we have implemented an easy way to transition between these two visualizations as described in the next sections. In VR mode an interactive visualization is presented to the user, allowing them to manipulate and visualize single-cell data using an array of annotations through the cardboard visor. Additionally, *singlecellVR* features the ability to receive as inputs, the standard output files of commonly-used tools for standard single-cell analysis: *Seurat* (Hao et al., 2021), *Scanpy (along with EpiScanpy)* (Wolf et al., 2018; Danese et al., 2019), *STREAM* (Chen et al., 2019a), *PAGA* (Wolf et al., 2019), and *scVelo* (Bergen et al., 2020). A companion package, *scvr* enables the conversion of these standard outputs to VR-compatible objects in a single command.

In the sections below, we will describe the basis for this VR visualization platform as well as provide descriptive examples of the visualization that can be performed using *singlecellVR*. We will compare *singlecellVR* to existing methods and describe its unique advantages that build on the early progress of single-cell data visualization in VR. We include a detailed protocol and quick-start guide that describe how the web platform enables researchers to explore their own data and dually functions as a database for preformatted datasets that can be explored immediately in VR (**Supplementary Note 1**).

### 2.2 VR Database and scvr preprocessing tool

*singlecellVR* provides a growing database of several datasets processed for VR visualization. Initialization and future growth of this database is enabled, scale-free through the streamlined *scvr* utility. As shown in **Figure 1**, to use *singlecellVR*, the user may select a precomputed dataset or convert their data from commonly used single-cell workflows. This conversion can be easily accomplished by using *scvr*, a simple one-line command tool for performing data conversion and produces a simple zipped .JSON file with all of the information required for visualizing cells and their annotations in VR. Additionally, datasets for which RNA velocity information has been calculated may be submitted directly for visualization of velocity in VR without prior conversion (**Supplementary Note 2, Supplementary Notebook 4**).

**Figure 1.**
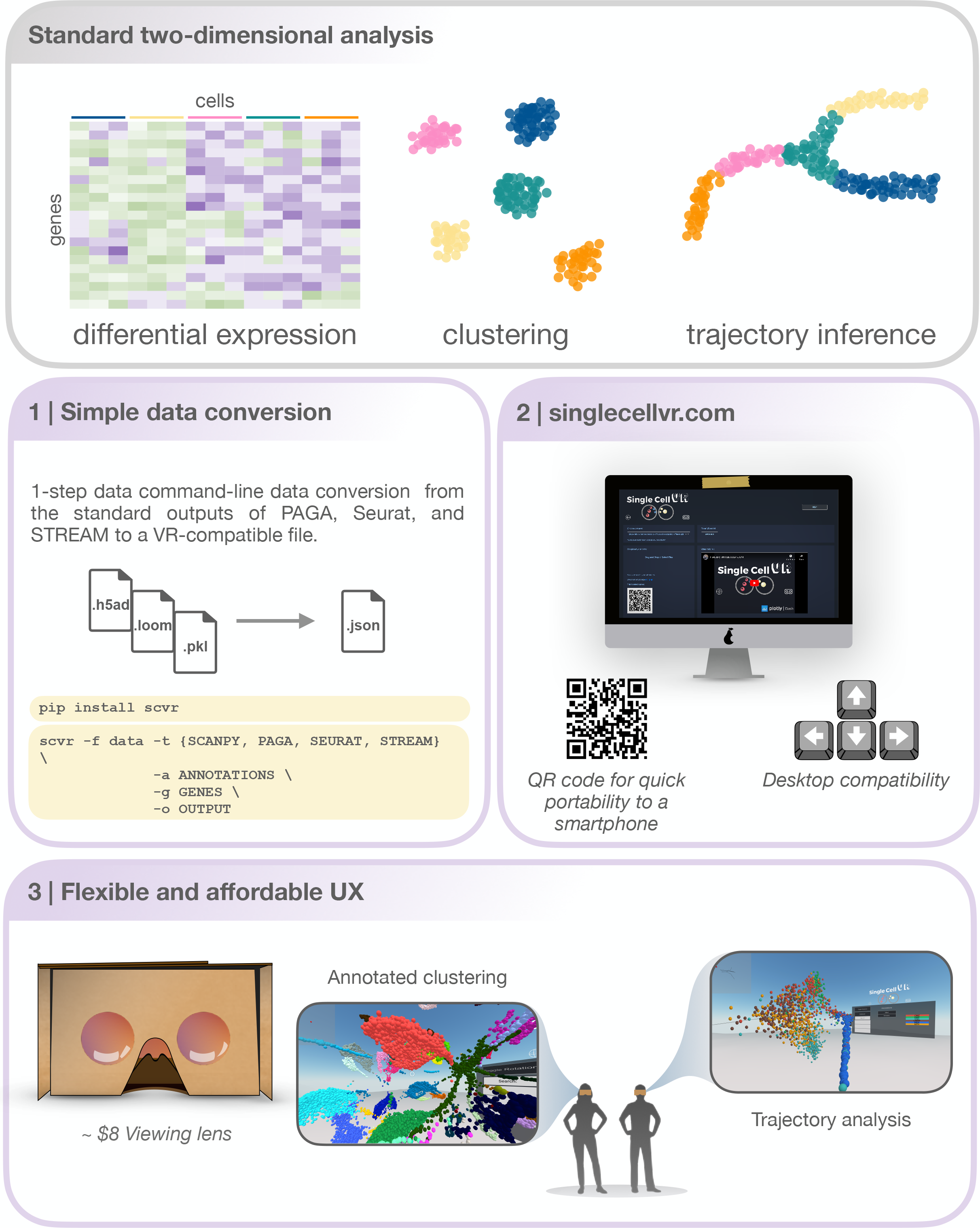
An overview of the *singlecellVR* user experience. Top, grey: The outputs of a standard 2-dimensional scRNA-seq analysis. Middle and bottom, purple: a step-by-step overview of the *singlecellVR* workflow: **1**. Schematic of *flexible* data conversion. One command to install (via the Python pip package manager) and one command to convert the data to be VR-compatible. **2**. Webpage for uploading and exploring VR data. **3**. VR mode visualization using a cheap smartphone enabled headset.

Conversion from the standard output of any single-cell analysis tool to this format would normally pose a significant methodological roadblock to most users, especially non-computational biologists. To bridge this gap, *scvr* parses and converts the outputs of *Scanpy, EpiScanpy, Seurat, PAGA*, and *STREAM* (respectively .*loom*, .*h5ad* and .*pkl*) and creates the required zipped .*JSON* file (Supplementary Note 2). This file contains the 3-D coordinates of cells in a specified space (e.g. UMAP, LLE, etc.), cell annotations (e.g. FACS-sorting labels, clustering solutions, sampling time or pseudotime, etc.), and feature quantification (gene expression levels, transcription factor deviation, etc.). It also contains the graph structure (the coordinates of nodes and edges) obtained from supported trajectory inference methods. Users interested in visualizing scRNA-seq dynamics using RNA velocity generated from spliced and un-spliced read counts can likewise prepare this information for visualization in *singlecellVR* using the *scvr* companion utility. Users can follow established workflows for obtaining these insights from the raw read file inputs as well as make use of the tutorials available at the *singlecellVR* GitHub Repository (**Materials and Methods, Supplementary Notebook 4**).

Importantly, *scvr* has been made available as a Python pip package to streamline its installation and can convert a processed dataset for VR visualization with a simple command. To install, one can simply open their command line utility and run: “pip install scvr”. Once installation is completed, the user can navigate to https://github.com/pinellolab/singlecellvr, to copy and customize the example commands provided to execute the one-step process for converting their data to a VR-compatible format. In addition to the documentation of *scvr* we have filmed a short video tutorial found on the homepage of *singlecellVR* to further assist less experienced users in preparing their data for visualization.

To showcase the functionality and generalizability of *scvr* across data types, we have preprocessed a collection of 17 published datasets, which includes both scRNA-seq as well as scATAC-seq and single cell proteomic data and made them available for immediate VR visualization. Taken together we believe this step addresses a key limitations of previously-developed VR tools mentioned above, and a formal comparison is presented below (Yang et al., 2018; Legetth et al., 2019; Bressan et al., 2021).

Excitingly, given the small footprint of the files obtained with *scvr*, we are offering users the ability to easily submit their processed data to the *singlecellVR* GitHub Repository (see **Supplementary Figure 1**) to make the tool a general resource for the field. In this way, we hope to even further extend the ability of biologists to visualize once static datasets and easily generate new hypotheses through manipulation of a large number of rich datasets. Therefore, we envision that our website will function as a repository for VR visualization data of single cell biological annotations.

### 2.3 A simple, cloud-based web tool for VR visualization

*singlecellVR* is available as a webapp at https://www.singlecellvr.com. This website enables users to explore several pre-loaded datasets or upload their own datasets for VR visualization. To build *singlecellVR* we have adopted recent web technologies, *Dash* by *Plotly* and *A-FRAME*, a recently-developed JavaScript framework for VR/AR. This allowed us to create a tool that is portable and does not require any installation. The website can be reached through any web browser and browser compatibility was tested against Google Chrome, Apple Safari, and Mozilla Firefox. Visualization can be done either on a personal computer or smartphone (both Android and Apple smartphones).

Once the users have uploaded their data to *singlecellVR*, they have the option to view and explore the data in 3-D directly in their web browser or to quickly jettison the data to their mobile device for visualization in a VR headset (**Figure 2** and **Supplementary Figure 2**). A key challenge associated with developing a method for visualization of single-cell data is transporting data that is typically processed on desktop setting to the smartphone-based VR visualization. In fact, we predict that in most cases, users will prefer to upload their data through a computer in which they may have run their analyses. To overcome this challenge and enable a seamless transition to a smartphone for VR view, our website dynamically generates a QR code that enables users to open the VR view on their phone to view data uploaded through a personal computer. This mixed approach is particularly useful because, as mentioned before most users are not processing single-cell data analysis from a phone nor would they keep the files on a mobile device.

**Figure 2.**
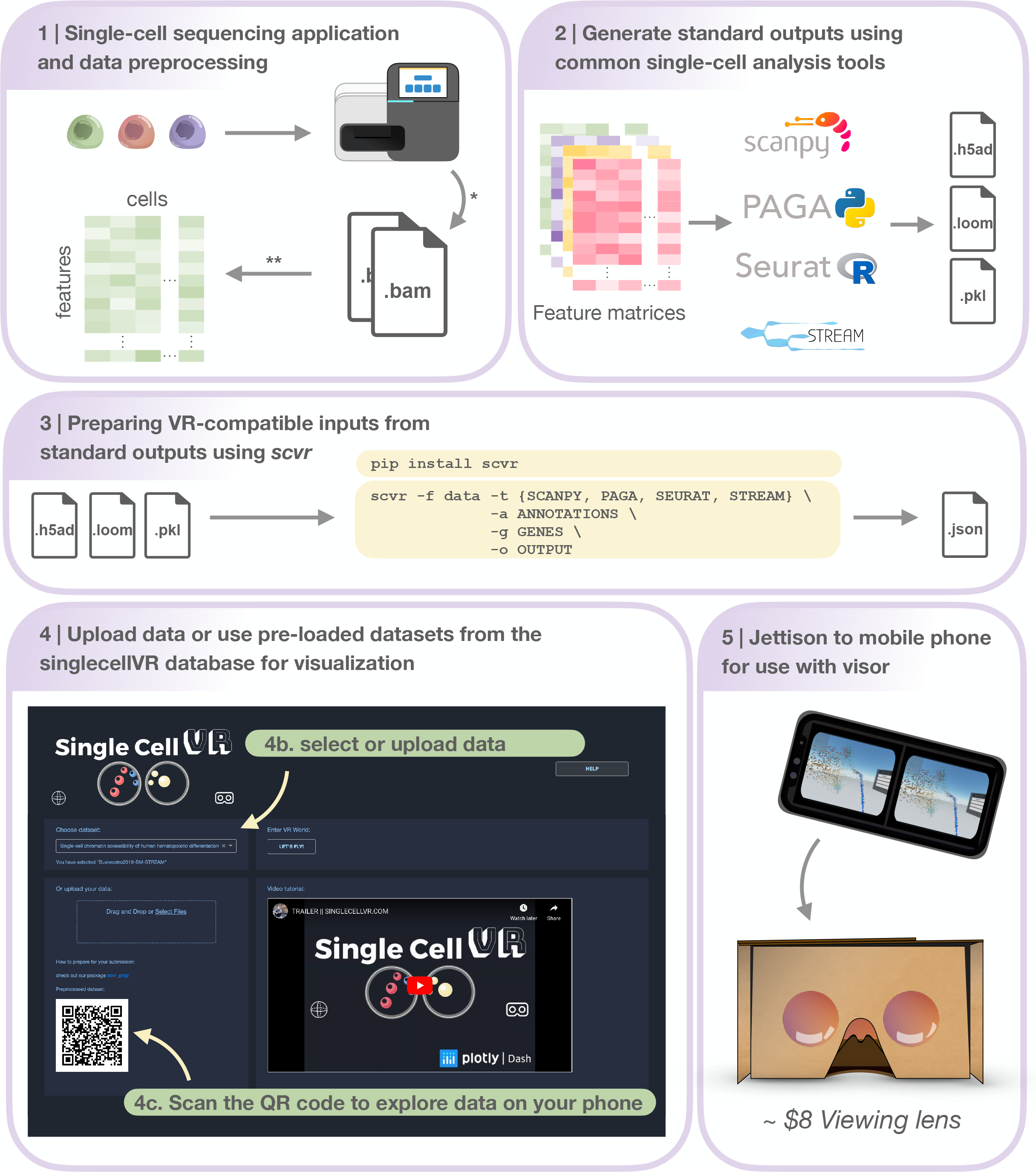
Step-by-step protocol for data processing and using *singlecellVR*. **Step 1**. Single-cell data can be generated using a variety of technologies or downloaded from online repositories*. Data can then be preprocessed and prepared for use (most often as a feature matrix) with common single-cell analysis tools**. **Step 2**. Pre-processed data can be analyzed using common single-cell analysis tools (listed here). **Step 3**. Users can process their data for use with *singlecellVR* from any of the standard outputs created by analysis tools listed in ***Step 2***. listed at the top in a single command. **Step 4**. Users can select from pre-processed data or upload their own data (***Step 4b***) and scan the dynamically generated QR code with their phone to begin the VR visualization (***Step 4c***). **Step 5**. Users can use the QR code on the website to transfer their data to their phone for use with simple hardware. * and ** are explained in the **Materials and Methods** section, ***Step 1***.

### 2.4 Supported tools and analysis

#### 2.4.1 Visualizing single-cell clustering solutions in VR

As previously mentioned, *Scanpy* and *Seurat* are two commonly-used tools for performing cell clustering as well as differential expression analysis. Here we demonstrate the utility of *singlecellVR* to visualize the common outputs of these tools, showcasing both the clustering solutions as well as differentially expressed genes or other technical or biological features that are visualized easily through the VR interface (**Figure 3**). A key advantage of our tool is the ability to supply multiple annotations to cells to visualize various attributes of the measured data, for example based on a biological query of interest or experimental design. This may include stratification by cluster identity, time points, tissues, or FACS-based labels. In **Figure 3**, we demonstrate the ability to select visualizations by various cluster identifications, which are user-customizable. With the advent of cross-experiment integration methods that can integrate not only multiple scRNA-seq experiments but experiments across modalities of single-cell data collection, this flexible labelling strategy should enable the user in the future to visualize even the most novel and complicated experiments in rich detail.

**Figure 3.**
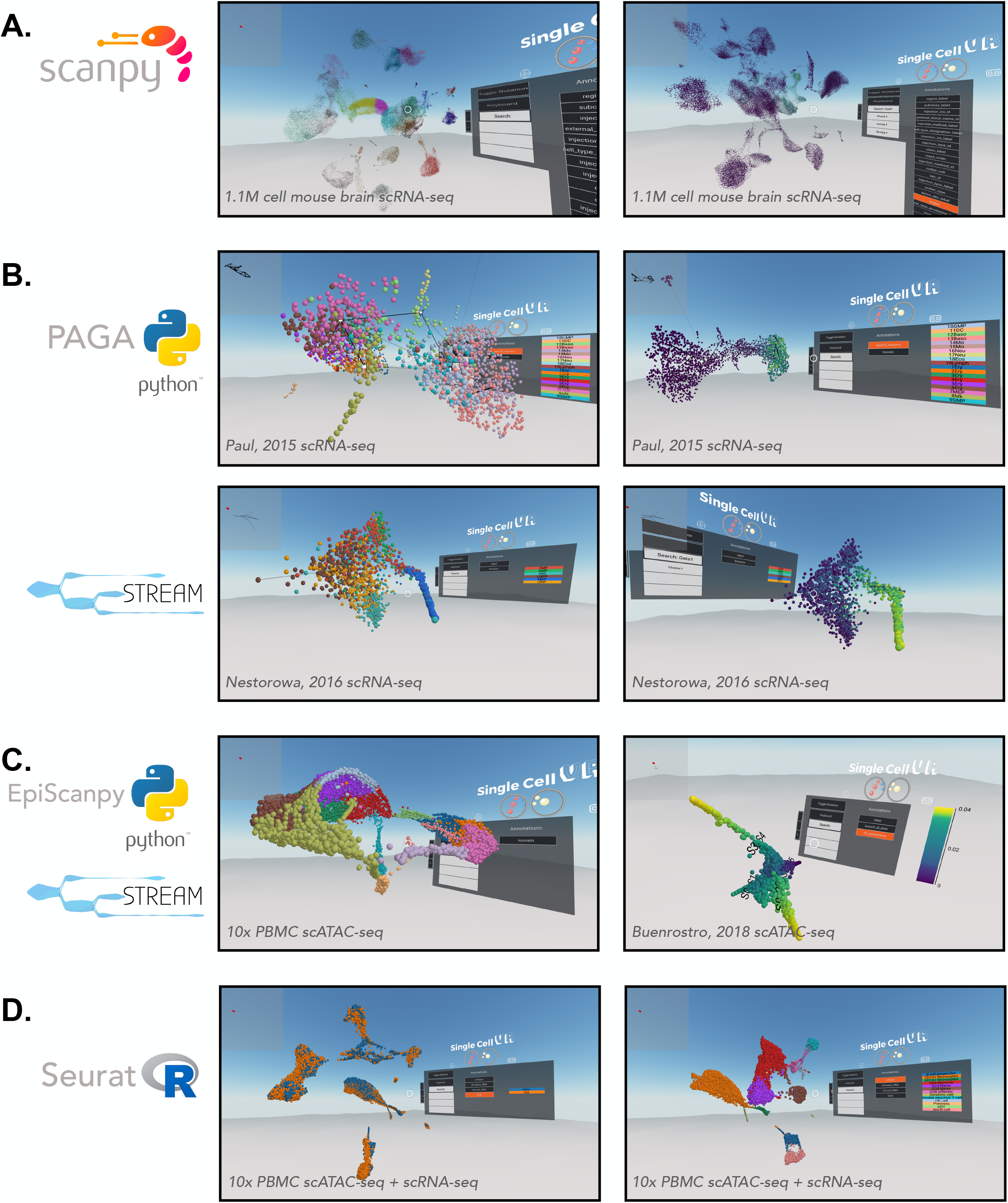
VR visualization of single-cell processed datasets profiled by different technologies and analyzed by various computational tools. **A**. Scanpy offers solutions for clustering single-cell data. Shown is a UMAP of the Allen Brain Atlas mouse brain scRNA-seq dataset from *Yao, et al*., 2020 (Yao, 2020) and processed by Scanpy. Leiden clustering solution (**left**) and expression of *Gad1* (**right**). **B**. Trajectory inference applications. *PAGA* offers a partition-based graph abstraction to uncover potential trajectories (edges) between group of cells (nodes) (**top-left**) relative gene expression (e.g., *Klf1*, ***top-right***), amongst other annotations. The *PAGA*-analyzed dataset shown here is from *Paul, et al*., 2015 (Paul et al., 2015). *STREAM* offers the visualization of developmental trajectories, which can be visualized by cell identity (**bottom-left**) or by relative gene expression (e.g., *Gata1*, ***bottom-right***), amongst other annotations. The STREAM-analyzed dataset shown here is from *Nestorowa, et al*., 2016 (Nestorowa et al., 2016). **C**. Epigenomic applications. *EpiScanpy* enables the clustering and visualization of scATAC-seq data (**left**). PBMC (healthy donor) 10,000 cells dataset analyzed by *EpiScanpy* and with colors corresponding to clustering solutions (Louvain clustering). *STREAM* was used to perform trajectory inference on the *Buenrostro, et al*., 2018 scATAC-seq dataset (Buenrostro et al., 2018) (**right**). **D**. Seurat offers solutions for clustering single-cell data as well as integrating datasets across experiments. Shown is a Seurat-integrated scRNA-seq and scATAC-seq PBMC dataset from 10x Genomics, colored by technology (**left**) and cell type (**right**).

In addition to flexibility for visualizing complex experimental setups, *singlecellVR* is able to visualize large experiments. To demonstrate this utility, we first processed (using *Scanpy* and *scvr*) and visualized on *singlecellVR*, scRNA-seq data from the Chan-Zuckerberg Biohub *Tabula Muris* project, a dataset consisting of 44,949 cells and 20 tissues from seven mice (Schaum et al., 2018). In **Supplementary Figure 3A**, clustering analyses of this dataset are projected into VR, colored by mouse tissue (**left**) and Louvain cluster identity (**right**). With a quick rendering time (<1 second) for the *Tabula Muris* dataset, we next explored the realm of visualization for a modern, large atlas-scale dataset (>1M cells). Using *Scanpy* and *scvr*, we successfully processed and visualized on our website, cells from the Allen Brain Institute that capture cortical and hippocampal development inside the mouse brain (**Figure 3A**) (Yao, 2020). This dataset consists of 1,093,785 cells and is among the largest scRNA-seq datasets created, to date. Visualization of this dataset in a dynamic VR setting creates the opportunity for more in-depth study of sub-sections of the data, which is particularly valuable for such a large dataset.

#### 2.4.2 Visualizing single-cell trajectory inference results in VR

Single-cell measurements are particularly useful for capturing cross-section snapshots of a biological process. With dense cell sampling, one can often observe transient cell states that exist between two, more stable states. However, without an intrinsic understanding of the process being studied, it may be difficult to order these cells along a time axis of a biological process. To enable ordering cells by transcriptional (or epigenomic) states, pseudotemporal ordering, based on trajectory inference and machine learning algorithms has become a useful technique for the single-cell field. Trajectory inference, like clustering, describes a high-dimensional biological process and being limited to a two/three-dimensional static visualization on paper, with a limited selection of genes or annotations is not ideal. Thus, we intend for our tool to leverage the richness of these datasets and make their general usefulness to the field more widespread. We therefore wanted to extend our VR visualization to the results of common trajectory inference tools (**Figure 3B**). *singlecellVR* supports two trajectory inference tools: *PAGA*, a partition-based graph abstraction trajectory inference method and *STREAM*, a method based on principal graphs (Albergante et al., 2020) that recovers a tree like structure to summarize developmental trajectories and to visualize the relative densities of cell populations along each branch.

To showcase the ability of *singlecellVR* to visualize trajectory inference results, we reprocessed a popular myeloid and erythroid differentiation dataset (Paul et al., 2015), performing trajectory inference using *PAGA. PAGA* is designed specifically to preserve relative cell topology in constructing the trajectory along a pseudotime axis. In the depiction of the *PAGA*-generated trajectory, nodes (gray) correspond to cell groups, and edges (black lines between nodes) connecting the groups quantify their connectivity and confidence (thickness) (**Figure 3B, top**). To showcase the VR output of *STREAM* we reprocessed a popular mouse blood dataset (Nestorowa et al., 2016). In STREAM, a set of smooth curves, termed *principal graph*, are fitted to the data and each curve represents a developmental branch. Within *singlecellVR*, we are able to easily explore these trajectories and observe qualitatively, the distribution of cells along each branch in the UMAP space (**Figure 3B, bottom**). The branches of these trajectories are represented by the curves that cut through the cells.

*singlecellVR* and *scvr* also support processing and visualizing single-cell epigenomic data. To demonstrate this functionality, we first used the *EpiScanpy* workflow to cluster a scATAC-seq dataset from 10x Genomics containing 10,000 cell PBMC (healthy donor) (**Figure 3C, left**). Next we reprocessed with STREAM a scATAC-seq dataset profiling human hematopoiesis (Buenrostro et al., 2018) (**Figure 3C, right**). In addition, we extend *singlecellVR* to single-cell quantitative proteomics data. To this end we reprocessed data from *SCoPE2*, a recent assay to quantitate proteins in single cells using mass spectrometry (Specht et al., 2019). We performed trajectory inference using *STREAM* on one *SCoPE2* dataset profiling the transition from monocytes to macrophages in the absence of polarizing cytokines. Our analysis revealed a bifurcated branch structure as cells progress towards macrophage phenotypes (**Supplemental Figure 3B**). Importantly, such bifurcation is not readily visualized in previous reports in two dimensions. Finally, we took advantage of the recent advances in the multi-omics field, using *Seurat* to integrate and co-embed PBMC cells profiled by scRNA-seq and scATAC-seq by *10x Genomics* (**Figure 3D**).

#### 2.4.3 Visualizing RNA Velocity Analysis in VR

Having successfully applied the *singlecellVR* framework to the visualization of trajectory inference analyses for multiple modalities of single-cell data, we sought to extend the framework further to visualize dynamical changes at single-cell resolution by way of RNA velocity. We first demonstrated this on a popular endocrine pancreas dataset (**Figure 4**), which has been previously employed to demonstrate the utility of visualizing dynamic processes using velocity.

**Figure 4.**
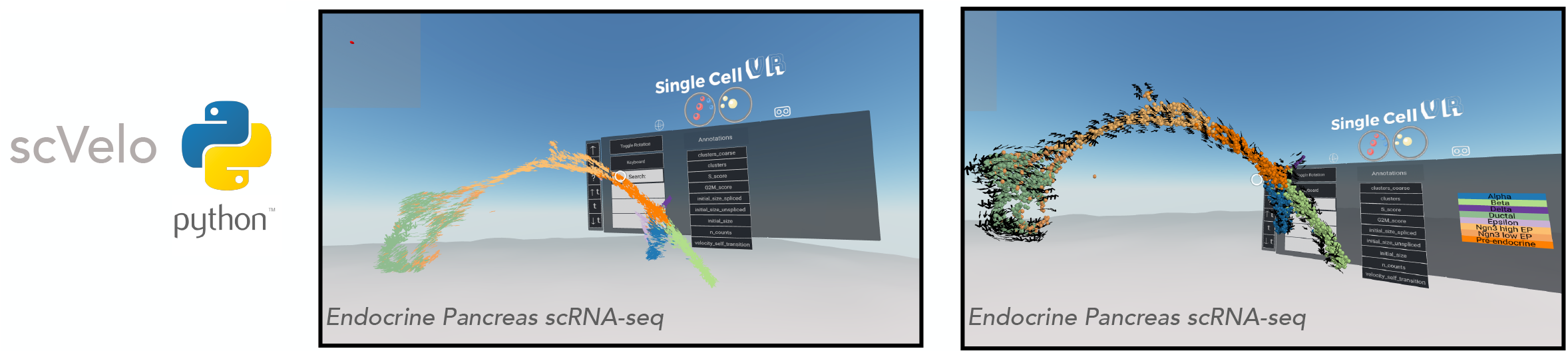
VR visualization of single-cell datasets with RNA Velocity. *scVelo* enables efficient analysis of the RNA velocity attributes of single-cell data. Shown is a 3-D UMAP of an endocrine pancreas dataset (Bastidas-Ponce et al., 2019). **Left**: Cells are displayed as their corresponding 3-D velocity vectors and colored according to cluster annotation. **Right**: Cells are displayed as 3-D orbs surrounded by a corresponding grid of velocity vectors. Cells are colored according to cluster annotation.

Visualization of RNA velocity using *singlecellVR* has two modes. In the default mode, each cell is represented by an arrow where the magnitude and direction of the arrow denote the velocity of that cell (**Figure 4, left**). For larger datasets, cells may be represented as spheres while a surrounding grid system of arrows denotes the predicted trajectory of a given cell (**Figure 4, right**). This is particularly helpful for interpreting the overall direction of cells in various clustering regions or subsets of a given trajectory. In either mode, the arrows are animated to gravitate towards the direction of the corresponding cell trajectory. Latent time, *t* is a parameter of the velocity calculation for a given cell. To aid in user comprehension of observed velocity, the speed and distance of the animated velocity vector may be calibrated on the fly during the VR experience through adjustment of the *t* parameter using the floating VR assistance menu. These results taken together with the visualizations of clustering analyses as well as trajectory inference analyses indicate that *singlecellVR* is a robust, generalizable tool across multiple modalities of single-cell analysis.

### 2.5 Comparison of singlecellVR to existing methods

As mentioned above, there are currently three unpublished reports of VR tools created to visualize single-cell data: *CellexalVR* (Legetth et al., 2019), *starmap* (Yang et al., 2018), and *Theia* (Bressan et al., 2021). In this section, we compare these tools to *singlecellVR* on two axes: (1) ease of use and (2) overall performance for visualization and analysis in VR.

#### 2.5.1 Ease of use

##### 2.5.1.a competing tools

Both *CellexalVR* and *Theia* require or recommend *HTC Vive* or *HTC Vive Pro* VR hardware (∼$500-1000), an *Intel Core i7 processor* (∼$300) or better, *an NVIDIA GTX1080* or *NVIDIA GeForce RTX 3080/3090 (∼$1500-3000), 16-32 GB RAM (∼$50-150) and a solid-state hard drive (SSD) (1 TB SSD recommended for CellexalVR*) (∼$50-100). Altogether, this equipment requires a minimum investment of roughly $2300-$4550. These are computational equipment that most biologists will not have at their disposal within their lab, likely limiting use of this tool to more computationally-focused labs.

*CellexalVR* requires software and boilerplate-level to pre-process the data in preparation for VR visualization is required and therefore requires the user to perform scripting to prepare data for downstream use with the VR visualization. While *Theia* has provided a convenient python script to convert *AnnData* objects, their software is not open-source, hindering further community contribution.

A contrasting alternative to *CellexalVR* and *Theia* is *Starmap*, which is compatible with low-cost hardware such as *Google Cardboard*. However, *Starmap* takes as input comma-separated values containing information of the three-dimensional coordinates of cells in the visualization as well as annotations (e.g., cluster ID), and up to 12 features per cell. This file must be prepared entirely by the user without assistance from the *Starmap* platform, limiting the audience of this tool to experienced computational biologists.

##### 2.5.1.b singlecellVR

The single-command companion package for data preparation, *scvr* described above enables users to visualize their own precomputed data directly from the outputs of commonly-used single-cell RNA-seq analysis tools. Currently supported tools include *Scanpy, EpiScanpy, Seurat, PAGA, STREAM, and scVelo. singlecellVR* is the only tool of the three discussed that features a QR code to quickly transport the VR data visualization to another device.

#### 2.5.2 VR performance and analysis capabilities

##### 2.5.2.a competing tools

*CellexalVR* proposes a versatile, user-friendly visualization for standard scRNA-seq workflow outputs and demonstrates comparable utility on scATAC-seq data. *Theia* offers a similarly high-performance visualization of single-cell data; their key distinguishing contribution is their visualization of spatial transcriptomic single-cell datasets.

*Starmap* is only demonstrated on scRNA-seq data and lacks the ability to visualize analyses beyond clustering (such as trajectory inference or an illustration of velocity). Further, *Starmap* is only capable of displaying up to 12 features for a given cell, limiting the throughput with which users may analyze their data.

##### 2.5.2.b singlecellVR

In contrast to existing methods, *singlecellVR* offers both a high-performance visualization with in-depth analysis and the ability to visualize all modalities of data at scale, while at the same time offering a software that is compatible with low-cost hardware and requires minimal computational abilities. These advances, which build on the progress made by these initial methods create a tool that matches and, in some cases, exceeds the capabilities of the highest-performing available tools as well as flexibility of input modality with virtually zero barrier to entry.

## 3 Discussion

The amount of publicly available scRNA-seq data has exploded in recent years. With new assays to capture chromatin accessibility, DNA methylation and protein levels in single cells, we predict a second wave of dataset generation. Each of these datasets are extremely high-dimensional and thus, rich with latent information about a given biological sample. Ideally, biologists would be able to explore this treasure-trove of data from any angle and make hypotheses assisted by *in silico* analysis at little to no time cost. Often however, experimental biologists lack the advanced computational skills and/or time required to reprocess and reanalyze raw data from published experiments to gain an understanding of the data from their desired angle of interest. Additionally, biologists who wish to thoroughly explore data prior to publication may rely on a computational specialist who is less connected to the biological problem of interest, introducing a disconnect in hypothesis-driven experimental turnover.

While once primarily reserved for entertainment, VR has found utility in both industrial and academic applications. In this manuscript we present a protocol for visualizing single-cell data in VR. This protocol is based on *singlecellVR* a VR-based visualization platform for single cell data and discuss its innovations and differences with existing methods. Importantly, we provide a simple mechanism to prepare results from commonly-used single-cell analysis tools for VR visualization with a single command to considerably increase accessibility (see **Materials and Methods**). With this added utility, we seek to empower non-computational biologists to explore their data and employ rapid hypothesis testing that could not be made from the traditional static representations typical of communication in a scientific report on paper or a computer screen.

We anticipate that VR will become increasingly useful as a research and education tool and that the construction of software libraries will aid such advancements. VR has also recently found application in other sources of biological data, including single-neuron morphological imaging data (Wang et al., 2019), three-dimensional confocal microscopy data for fluorescent molecule localization (i.e., fluorophore-tagged proteins) within cells (Stefani et al., 2018), and three-dimensional single-molecule localization super-resolution microscopy (Spark et al., 2020). Our scalable and flexible VR visualization framework is not limited to scRNA-seq and it can be also easily adapted to other single-cell assays and tools that already support epigenomic data and/or single-cell proteomic data (*EpiScanpy* (Danese et al., 2019), *Seurat* (Stuart et al., 2019), and *STREAM* (Chen et al., 2019b)). Finally, we extend our framework to computational methods that derive the RNA velocity of single cells for visualization in VR(La Manno et al., 2018; Bergen et al., 2020). With the recent advances in spatially-resolved transcriptomics (Welch et al., 2019) and corresponding analysis methods (Hao et al., 2021; Miller et al., 2021), visualization of such data has already been extended to a VR framework (Bressan et al., 2021). We believe this new sort of three-dimensional VR will also become especially useful once made available to the general research community via inexpensive hardware and facile data preprocessing and preparation for VR visualization. As software to analyze single cells reach their maturity, one could imagine the incorporation of such visualizations into more clinically translatable settings, such as medical devices.

## 4 Conclusion

This manuscript presents *singlecellVR, a scalable* web platform for the VR visualization of sinlge-cell data and its associated preprocessing software, *scvr*, which streamlines the results of commonly used single-cell workflows for visualization in VR. *singlecellVR* enables any researcher to easily visualize single-cell data in VR. The platform is user-friendly, requires no advanced technical skills or dedicated hardware. Importantly, we have curated and preprocessed several recent single-cell datasets from key studies across various modalities of data generation and analysis approaches, providing the scientific community with an important resource from which they may readily explore and extract biological insight.

### KEY POINTS

- *singlecellVR* is a web platform that enables quick and easy visualization of single-cell data in virtual reality. This is highlighted by a database of pre-loaded datasets ready for exploration at a single click or via a QR code to quickly jettison the visualization to a smartphone enabled VR visor.
- *scvr* is a companion package to easily convert standard outputs of common single-cell tools in a single command
- *singlecellVR* is made for use with cheap and easily-available VR hardware such as Google Cardboard (∼ $10).
- *singlecellVR* can visualize both clustering solutions as well as trajectory inference models of single-cell data for transcriptomic, epigenomic, and proteomic data as well as multi-modally integrated datasets. Additionally, *singlecellVR* offers a three-dimensional VR visualization of RNA velocity dynamics.

## 5 Materials and Methods

### Single-cell data preparation

All datasets were processed using *Scanpy* (version 1.5.1, RRID:SCR_018139), *AnnData* (verson 0.7.6, RRID:SCR_018209), *EpiScanpy* (version 0.1.8), *Seurat* (version 3.1.5, RRID:SCR_007322), *PAGA* (part of *Scanpy*, version 1.5.1, RRID:SCR_018139), *STREAM* (version 1.0), and *scVelo* (version 0.2.3, RRID:SCR_018168) following their documentations. Jupyter notebooks to reproduce data processing are available at https://github.com/pinellolab/singlecellvr. Analyses were performed on a 2019 MacBook Pro (2.4 GHz Intel Core i9, 16 GB RAM).

### Preparation of processed data for visualization in VR

The preprocessing package, *scvr* generates a series of .json files containing the spatial coordinates representative of cell embeddings in 3D embedding (e.g. PCA, UMAP, etc.) and information including labels and features (e.g., gene expression, TF motif deviation, etc). These .json files are zipped upon output from *scvr* into a single file that can be easily uploaded to *singlecellVR* for visualization.

### singlecellVR webapp construction

To build *singlecellVR*, we used *A-FRAME* (version 1.2.0), *Dash* by *Plotly* (version 1.13.3).

## Supporting information

Supplementary IPython Notebooks

## 7 Author Contributions

DFS, HC, MEV, and LP conceived this project and designed the experiment, which was begun at the 2019 HackSeq, at the University of British Columbia where input from collaborators mentioned in the acknowledgements was received. DFS, HC, MEV, and RDC processed the data hosted in the database as well as produced the VR image demonstrations of *singlecellVR* shown in this manuscript. DFS led the development of the virtual reality framework. HC led the development of Dash-based website and the preprocessing module, scvr. QQ contributed to extending the VR tool with velocity and new API. QZ contributed to extending the VR tool to incorporate single-cell protein analysis. MEV led the preparation of the manuscript. All authors performed user-testing of the software. LP supervised the development of this work and provided guidance. All authors wrote and approved the final manuscript.

## 9 Acknowledgments

We would like to acknowledge the organizers and participants of *HackSeq19 (University of British Columbia)*, where this project began. We would like to especially acknowledge those *HackSeq19* participants that contributed to the development of this project: Michelle Crown, Alexander Dungate, David Lin, Terry Lin, and Sepand Dyanatkar Motaghed. Additionally, we would like to thank Ashley Browne, Salma Ibrahim, and Kate McCurley for their contributions to the construction of the *singlecellVR* database through the Winsor Science Internship Program.

## 10 Data Availability Statement

The dataset shown in **Figure 3A** is from *A taxonomy of transcriptomic cell types across the isocortex and hippocampal formation* 2020 (Yao, 2020) and was downloaded and processed from https://portal.brain-map.org/atlases-and-data/rnaseq/mouse-whole-cortex-and-hippocampus-10x. The dataset shown in **Figure 3B** (**top**) is from *Paul, et al*. 2015 (Paul et al., 2015) and was downloaded from GEO, accession: GSE72859. The dataset shown in **Figure 3B** (**bottom**) is from *A single-cell resolution map of mouse hematopoietic stem and progenitor cell differentiation* (Nestorowa et al., 2016) and was downloaded from GEO, accession: GSE81682. The scATAC dataset shown in the clustering result of **Figure 3C** (**left**) is the 10x PBMC (healthy donor) generated by 10x Genomics and was downloaded from here: https://support.10xgenomics.com/single-cell-atac/datasets/1.2.0/atac_pbmc_10k_v1. The scATAC dataset shown in the trajectory inference result in **Figure 3C** (**right**) is from *Integrated Single-Cell Analysis Maps the Continuous Regulatory Landscape of Human Hematopoietic Differentiation* (Buenrostro et al., 2018) and was downloaded from GEO, accession: GSE96769. The scRNA-seq and scATAC-seq data shown in **Figure 3D** was obtained from the *Seurat v3* vignette available here: https://satijalab.org/seurat/v3.2/atacseq_integration_vignette.html. The endocrine pancreas dataset shown in **Figure 4** was generated in *Comprehensive single cell mRNA profiling reveals a detailed roadmap for pancreatic endocrinogenesis* (Bastidas-Ponce et al., 2019) and analyzed from the processed .h5ad data through the *scv.datasets.pancreas* function in *scVelo*. The dataset shown in **Supplementary Figure 3A** is from the Chan Zuckerberg *Tabula Muris* project (Schaum et al., 2018) and was downloaded from: https://figshare.com/projects/Tabula_Muris_Transcriptomic_characterization_of_20_organs_and_tissues_from_Mus_musculus_at_single_cell_resolution/27733. The data shown in **Supplementary Figure 3B** was obtained from https://scope2.slavovlab.net/docs/data.

## 11 Code Availability Statement

The source code and the supporting data for this study are available online on GitHub at https://github.com/pinellolab/singlecellvr. The preprocessing package, *scvr* is included within that repository https://pypi.org/project/scvr/. The documentation for scvr is available here: https://github.com/pinellolab/singlecellvr. Video tutorials for learning about and running visualization experiments with *singlecellVR* (and using *scvr* to prepare the data) are available on YouTube, here: https://www.youtube.com/playlist?list=PLXqLNtGqlbeMaAuiBStnBzUNE6a-ULYx8.

All the analyses in this manuscript can be reproduced using the Jupyter notebooks available at https://github.com/pinellolab/singlecellvr. Additionally, we have provided a wiki within the same repository for a more detailed guide to reproducing results from the paper as they pertain to the supplementary materials.

## 12 Ethics approval and consent to participate

Ethics approval was not needed for the study.

## Figure legends

**Supplementary Figure 1.**
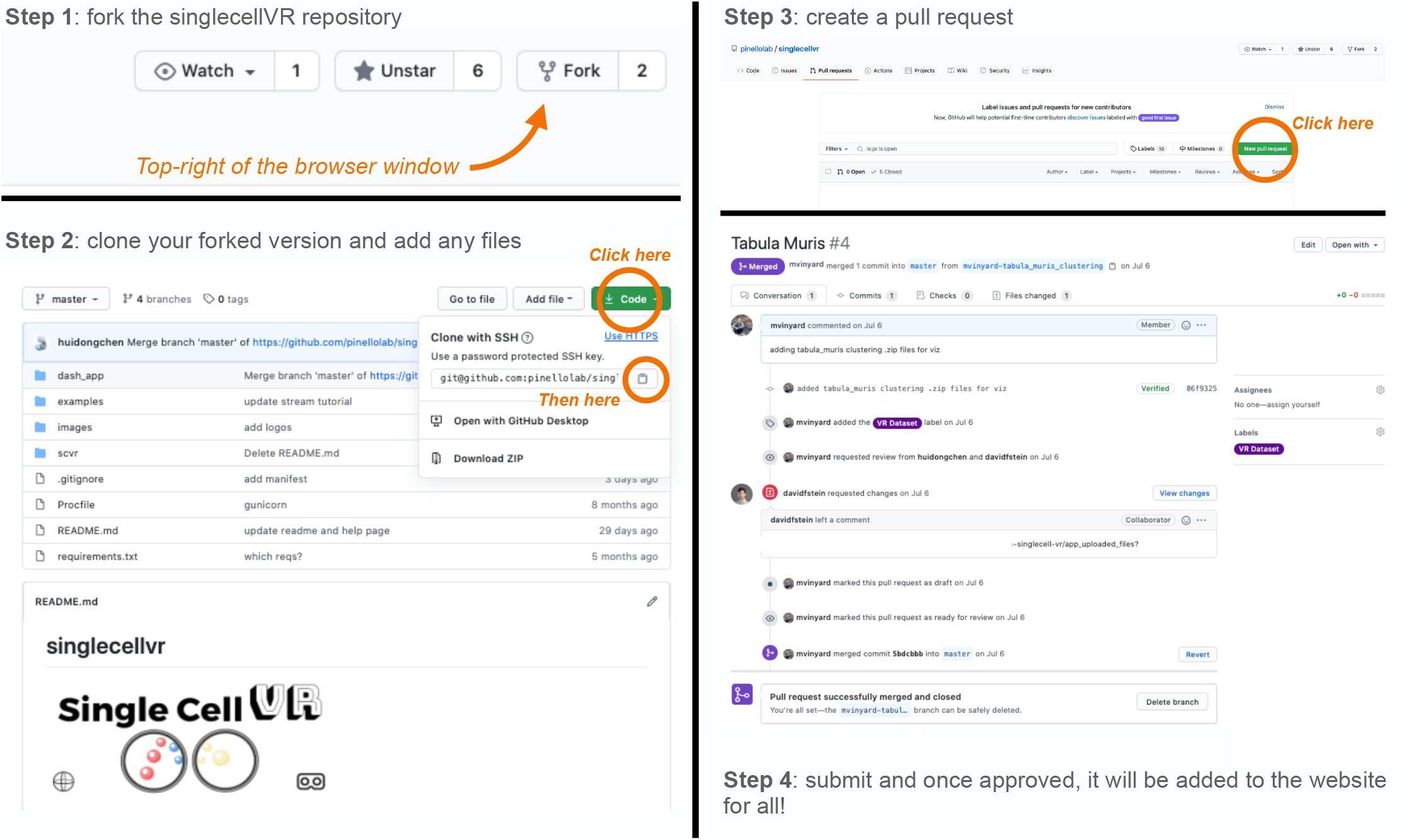
Instructions for contributing VR-processed data to the *singlecellVR* data repository. Users can contribute to the growing repository of VR datasets by submitting a pull request to our GitHub repository: https://github.com/pinellolab/singlecellvr. To do so, first fork and clone the repository (**steps 1** and **2**, above). Next, add your data (**step 3)**. Finally, create a pull request (**step 4**) to be submitted for approval. Once approved, your data will be incorporated into the growing repository of VR datasets. It is necessary to add the ‘VR Dataset’ flag (purple, already added to the sidebar) to the pull request. In addition, we ask users to describe the data, methods used and available annotations (e.g. genes, timepoints, clusters labels etc.) in the commit message or comment section of the pull request. ***Note***: *for velocity results, files >50 MB are too large to be shared through GitHub and must be shared via other channels. However, coordination of this sharing may proceed through GitHub as shown in this figure. For more, see* ***Supplementary Note 2*** *and* ***Supplementary Notebook 4***.

**Supplementary Figure 2.**
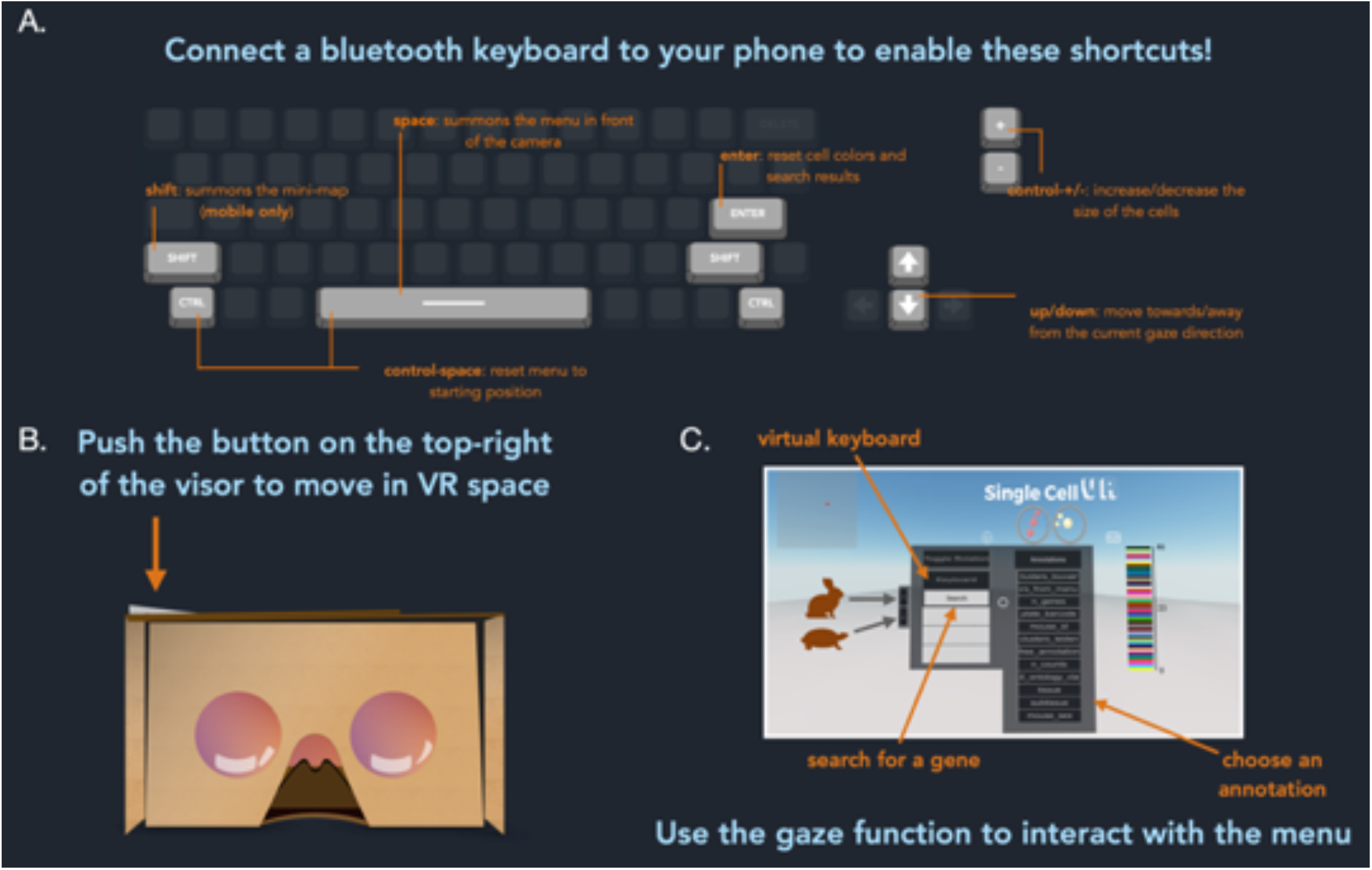
Tips for using the VR interface. **A**. It is not required but one can easily connect a keyboard with your smartphone using a Bluetooth-enabled keyboard (a small portable keyboard can be purchased from Amazon for ∼$10). However, you can still use a normal computer with your browser and explore using your mouse and keyboard, the three-dimensional transcriptional space with cells, trajectories and graph abstractions. The full set of interactive keyboard functionalities are detailed above. **B**. There are several similar versions of cardboard VR adapters available for ∼$10. Many VR headsets such as Google Cardboard have a single button that allows a user to click the screen of their phone while immersed in a virtual reality experience. By holding down this button, users without a keyboard may move forward in the direction of their gaze. You can also simply use your computer screen to do initial exploration of the data in 2-D. **C**. Users may navigate the VR visualization via a combination of gaze controls and keyboard inputs. A circle, centered in the user’s field of view indicates the direction that a user will move through the virtual space and also acts as the appendage through which the user will interact with objects in the visualization. Additionally, users may select the “keyboard” button on the menu to render a virtual keyboard. Cardboard users may use this keyboard to search for available features to render on the display. The “Enter/Return” key on the virtual keyboard clears the current search. Subsequently selecting the “keyboard” button will hide the keyboard from view.

**Supplemental Figure 3.**
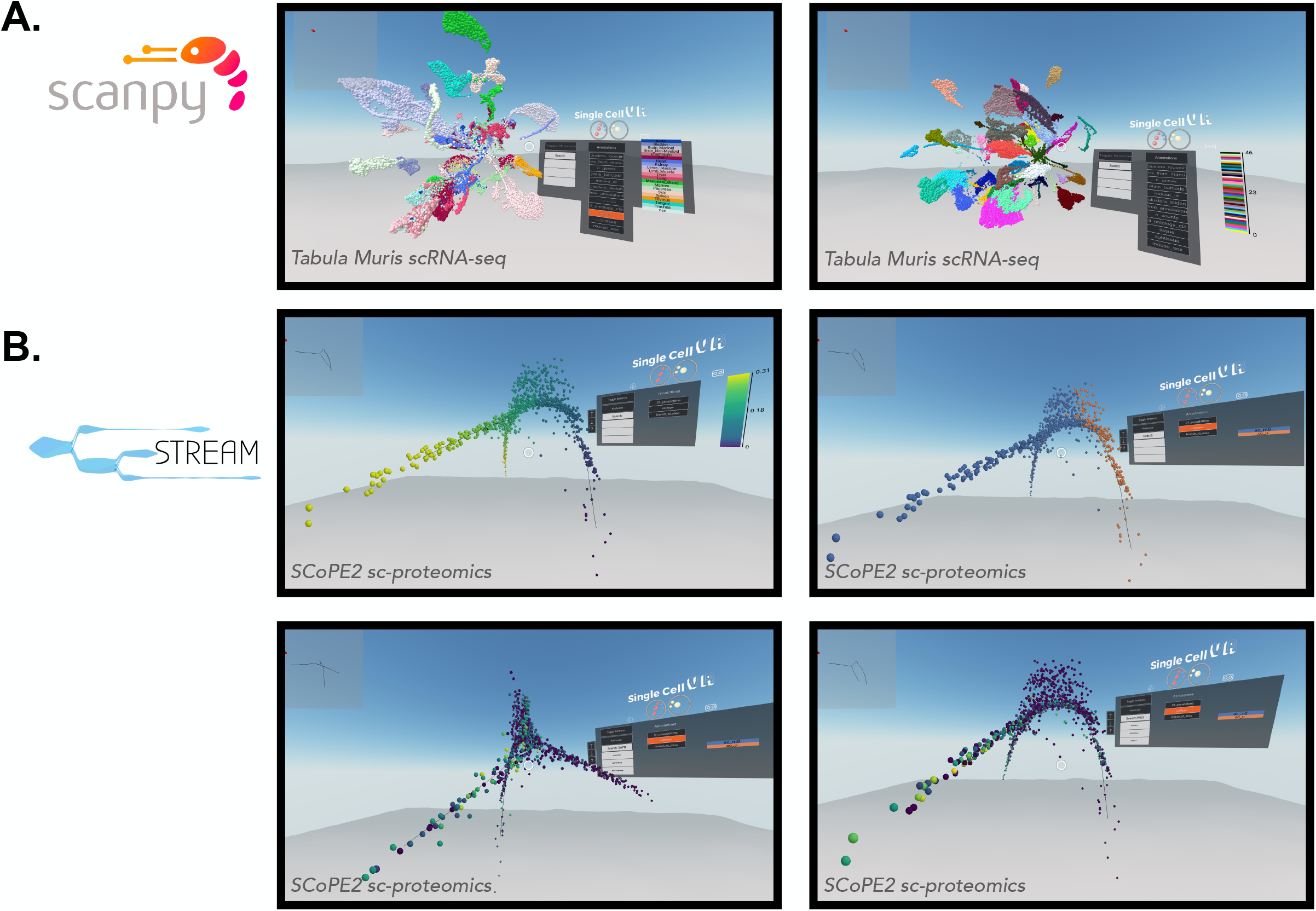
**A**. Rendering the single-cell virtual reality visualization. *Scanpy* offers tools for clustering, which can be visualized using *singlecellVR*. Cells can be visualized and colored by various annotations. Shown: mouse tissue type, (**left**) or their cluster ID (**right**). The *Scanpy*-analyzed dataset shown here is from the Chan Zuckerberg Initiative’s *Tabula Muris* dataset (Schaum et al., 2018). **B**. *STREAM*-processed single-cell proteomics data from *SCoPE2* (Specht et al., 2019). These visualizations are an example of an advantage gained by trajectory analysis and three-dimensional visualization. 3-D UMAP plots (ordered left to right, top to bottom) generated by *STREAM*, respectively colored by pseudotime progression, cell type (orange: monocyte, blue: macrophage), expression of *Safb*, and expression of *Pfn1*.

