## Supplementary Note 1 for "*singlecellVR*: interactive visualization of single-cell data in virtual reality"

**Supplementary Note 1**: Step-by-step instructions for preparing data for, and using singlecellVR

Here we summarize the capabilities of singlecellVR through a step-by-step protocol such that the user may learn how to use the tool the context of a typical data processing workflow. As such, we have included in each “step”, a detailed description of the required input and expected results. Therefore, for increased clarity, we have added a note to the end of each step indicating the expected file types that serve as input to a step and are subsequently created by that step. At the end of this note, there is a brief overview summary of all steps described.

**Step 1:** **obtain and preprocess data**.

An illustrative example for downloading and processing FASTQ files from public repositories is provided in **Supplementary Notebook 1**. This notebook details how to obtain the data with links to the sources as well as instructions for formatting and aligning the data to the genome using some of the most common tools.

***Input****: FASTQ files or processed data matrices*

***Output****: processed data matrices (cell x feature)*

**Step 2: processing your dataset with common single-cell analysis tools***.*

In short, we used several of the most common frameworks for post-processing and analysis of single-cell data: *Scanpy*, Seurat, Episcanpy, PAGA, and STREAM. We demonstrate the tool’s utility on scRNA-seq, scATAC-seq, paired scRNA-seq and scATAC-seq, scRNA-seq velocity, and sc-proteomic data.

For all of the single-cell analyses used to create the figures in this manuscript, we have provided a series of supplementary notebooks such that each dataset and the resulting visualization may be easily reproduced. The label of each notebook corresponds to the manuscript figure label in which the dataset is ultimately visualized in VR.

- **Supplementary Notebook 3A** details the *Scanpy* analysis of a 1.1 million cell mouse brain scRNA-seq dataset (shown in **Figure 3A**) (Wolf et al., 2018; Yao, 2020).
- **Supplementary Notebook 3B-1** offers the trajectory inference analysis of a myeloid and erythroid differentiation scRNA-seq dataset from Paul, et al., 2015 (Paul et al., 2015) using *PAGA* (corresponding to the visualization shown in **Figure 3B (top)**) (Wolf et al., 2019).
- **Supplementary Notebook 3B-2** details the *STREAM* trajectory inference analysis of a mouse blood scRNA-seq dataset from Nestorowa, et al., 2016 (Nestorowa et al., 2016; Chen et al., 2019) (corresponding to the visualization shown in **Figure 3B (bottom)**)**.**
- **Supplementary Notebook 3C-1** offers the *EpiScanpy* analysis of a scATAC-seq PBMC dataset from 10x Genomics (corresponding to the visualization shown in **Figure 3C (left)**) (Danese et al., 2019).
- **Supplementary Notebook 3C-2** details the *STREAM* analysis of a human hematopoietic differentiation scATAC-seq dataset from Buenrostro, et al., 2018 (Buenrostro et al., 2018; Chen et al., 2019) (corresponding to the visualization shown in **Figure 3C (right)**).
- **Supplementary Notebook 3D** shows the *Seurat* analysis used to analyze a paired scATAC-seq + scRNA-seq dataset from 10x Genomics (Stuart et al., 2019).
- **Supplementary Notebook 4** shows the *scVelo* analysis used to analyze data for which spliced and un-spliced reads had been analyzed to offer dynamical insights using *velocyto*(La Manno et al., 2018; Bergen et al., 2020).

***Input****: cell x feature matrix*

***Output****: data analysis object / file (.h5ad, .loom, .pkl)*

**Step 3: using the scvr preprocessing tool to prepare your data for VR**.

After easy installation using pip (**Step 3a**), the *scvr* command line tool runs with the execution of a single command from a terminal window.

Specifically, open a command-line terminal and then:

**Step 3a**. Install using `pip install scvr`

**Step 3b.** Once installation is completed, the user can navigate to <https://github.com/pinellolab/singlecellvr>, to copy and customize the example commands provided to execute the one-step process for converting their data to a VR-compatible format.

Scvr produces a relatively small file, which enables the continuous expansion of this VR-visualizable database. To submit your own data, please open a pull request at the *singlecellVR* GitHub page (<https://github.com/pinellolab/singlecellvr>).

***Input****: data analysis object / file (.h5ad, .loom, .pkl)*

***Output****: .JSON file compatible with singlecellVR*

**Step 4: uploading your data or selecting pre-loaded data for visualization in *singlecellVR***:

Users who have prepared their data using the *scvr* command-line tool described in **Step 3**, may upload their data to the website via a drag-and-drop interface. As described in the main text, a custom QR code is spontaneously generated, which enables easy portability to a mobile device for use with cheap smartphone-compatible hardware (e.g., Google Cardboard). Alternatively, users are able to view their data directly in the browser.

Using the *scvr* command-line tool described in **Step 3**, we have already preprocessed several datasets, including those which are shown in this manuscript. These preprocessed datasets are stored in a database configuration and may be loaded from a dropdown menu on the *singlecellVR* web interface such that users can get started with ease.

***Input****: .JSON file compatible with singlecellVR*

***Output****: QR code / visualization*

**Step 5: visualizing and analyzing single-cell data in VR**

Regardless of the analysis method from which their data was prepared, the user will here place their smartphone into the VR visor after scanning the QR code. Your data is now ready to be explored in VR – use your gaze (in VR) to interact with the menu and explore your data. For further assistance, we have produced a detailed walkthrough and users can use the tutorials published on YouTube, linked on the homepage of [https://singlecellvr.com](https://singlecellvr.com/).

***Step 5A****:* ***visualizing single-cell clustering solutions in VR****.*

The key user-interface to be aware of when visualizing clustering solutions in VR, are the menu options for toggling between various cluster labels and gene expression markers. The search bar enabled by an in-experience VR keyboard can be used to search for genes included in the dataset. Examples of visualizations using these results are featured in **Figure 3** and **Supplementary Figure 3**.

***Input****: Selected or uploaded dataset from Seurat or Scanpy, processed by scvr*

***Output****: Exploration of the cell subpopulations and their features*

***Step 5B****:* ***visualizing single-cell trajectory inference results in VR****.*

All functionalities enabled by the analysis of cluster-level visualizations in VR are available to the user when visualizing the *scvr*-converted outputs of single-cell trajectory inference methods. In addition, cells may be highlighted by attributes generated by these methods including pseudotemporal ordering. The key advantage of this combination of outputs with the VR framework is the ability to visualize the backbone structure of the 3-D trajectory, which can be observed within the cloud of cells. Examples of visualizations using these results are featured in **Figure 3** and **Supplementary Figure 3**.

***Input****: Selected or uploaded dataset from PAGA or STREAM*

***Output****: Exploration of the cell subpopulations and their features in VR*

***Step 5C****:* ***visualizing single-cell RNA velocity results in VR****.*

As in **Steps 5A** and **5B**, the user can search for genes and other cell annotations generated by *scVelo* and associated tools that may have deposited cell information in the predecessor *AnnData* object before *scvr* conversion. They key feature of the VR visualization focused on RNA Velocity is the ability to view the velocity vector for every cell (default, **Figure 4**, **left**). These vectors can also be project forwards in time by using the time-step enlargement or reduction buttons in the in-experience menu. Additionally, a grid of velocity vectors analogous to what one might observe while processing data in 2-D, using *scVelo* can be projected onto the 3-D VR visualization. This is especially useful for visualizing the overall dynamical movement of cell populations and subpopulations in the dimensionally-reduced projection space (**Figure 4**, **right**).

***Input****: Selected or uploaded dataset from scVelo*

***Output****: Exploration of the cell subpopulations and their features in VR*

Downstream preprocessing steps include filtering variable genes, k-nearest neighbors’ imputation, dimensionality reduction; these steps are performed using *scVelo*. Continuing with *scVelo*, a steady-state model of velocity is fit to each gene. “Velocity genes” are then determined through a best-fitting approach to the RNA velocity model. Non-velocity genes are filtered and the dimensionally reduced measurement is then projected onto a three-dimensional UMAP plot, relying on the assumption that linear expression changes along several constant time points wherein each next neighbor is a forward step in time and, for 0.1, 1, 10, or 100 steps may be taken to study the dynamics of a given cell. The start and end coordinates of these time steps are stored within an *AnnData* object.

**Quick-start reference**

This note, as a companion to the manuscript should serve to cover the major functional aspects of singlecellVR and the data preprocessing steps that proceed it. We have summarized the above steps as a quick-reference user-guide below:

*If using pre-loaded data, skip to* ***step 3****. If starting from unprocessed data, start with step 1.*

1. Download a dataset from public repositories and pre-process it using common pipelines to obtain a feature matrix.
2. Analyze the feature matrix with *Scanpy, Seurat, STREAM, or PAGA* to produce clusters, trajectories and biological relevant annotations.
3. **3a**. Install *scvr* by simply using “pip install scvr” (if you have already done this, there is no need to repeat this step and you can skip to the next step).

**3b**. Use *scvr* to convert your processed data (**Step 3**) to a VR-compatible format with the *singlecellVR* webapp (zipped .JSON file).

4. **4a**. Navigate to [https://singlecellvr.com](https://singlecellvr.com/)

**4b**. Click and drag your zipped VR-compatible results file into the indicated upload box *or* choose a pre-loaded dataset from the dropdown menu.

**4c**. Once the data has loaded, click *“let’s fly”* to begin your VR experience. If you have purchased the cardboard hardware, open the camera on your mobile phone and scan the QR code that is generated upon data loading. This will open the visualization in VR mode in a browser on your phone.

5. (**a**: clustering solutions, **b**: trajectory inference, **c**: RNA Velocity). Place your phone in the VR visor and explore your data. You can use your gaze (in VR) to interact with the menu and explore your data. For a detailed walkthrough, you can use the tutorials published on YouTube, linked on the homepage of [https://singlecellvr.com](https://singlecellvr.com/).
