## Supplementary Note 2 for "*singlecellVR*: interactive visualization of single-cell data in virtual reality"

**Supplementary Note 2**: **Preparation of scRNA-seq data for three-dimensional VR velocity visualization**

To process scRNA-seq data for visualization with *singlecellVR*, we first used *Velocyto* (version 0.1.7, RRID:SCR_018167) to extract spliced and un-spliced read counts for each gene or feature measured in each single cell within the experiment. This was performed using the unprocessed single-cell sequencing data – typically a BAM file or FASTQ file as input. The outputs of this method are a set of feature counts matrices (for both spliced and un-spliced reads) as well as other tabulated feature metrics generated from the counts data. The format of these outputs is typically a .*loom* and/or .*h5ad* file.

Downstream preprocessing steps include selecting top variable genes as well as performing *k*-nearest neighbors’ imputation and dimensionality reduction; these steps are performed using *scVelo*. Continuing with *scVelo*, an RNA velocity model is fitted to each gene. “Velocity genes” are then selected using a gene-fitting likelihood threshold of the RNA velocity model. The velocity genes’ spliced expression are then used to jointly project the current and future state of the cells onto a three-dimensional UMAP (3-D UMAP) plot, relying on the assumption that expression changes linearly along several constant time points wherein each next neighbor is a forward step in time. 0.1, 1, 10, or 100 steps may be taken to study the dynamics of a given cell. The initial and final 3D-UMAP coordinates of these time steps are stored within an *AnnData* object. Support for visualization may be initiated through GitHub (**Supplementary Figure 1**) though AnnData objects must be shared via separate channels due to the file size constraints of GitHub.
