## Supplementary IPython Notebooks for "*singlecellVR*: interactive visualization of single-cell data in virtual reality": Supplementary_Notebook_1.ipynb

```
# %% [markdown]
### Supplementary Notebook 1
---
### Standard preprocessing of single-cell data

#### Overview of steps performed:

1. **Download the data**


2. **Raw read alignment** (5 options, A-E)

    A. CellRanger
    
    B. STAR
    
    C. BWA
    
    D. Kallisto Bustools
    
    E. Salmon
    
    
3. **Feature matrix generation using HTseq-counts**

# %% [markdown]
### Step 1. Download the raw data

# %% [markdown]
#### Download data: 10x Genomics

Now, we have a few options. For the 10x Genomics data, the easiest thing to do is to use *CellRanger*, which produces a convenient assembly of files and provides QC metrics to assess the quality of the data. *CellRanger* also produces the count matrices in convenient file format. *CellRanger* uses *STAR* to align to the genome. This particular sample will be aligned to the `hg38` (downloaded <a href="https://cf.10xgenomics.com/supp/cell-exp/refdata-gex-GRCh38-2020-A.tar.gz">here</a>)

`curl -O https://cf.10xgenomics.com/supp/cell-exp/refdata-gex-GRCh38-2020-A.tar.gz`

`tar -zxvf refdata-gex-GRCh38-2020-A.tar.gz`

For this example, we will first start by downloading the data. We're using data from 10x Genomics: <a href="https://support.10xgenomics.com/single-cell-gene-expression/datasets/3.0.0/pbmc_1k_v3">5k Peripheral blood mononuclear cells (PBMCs)</a>

`wget https://cf.10xgenomics.com/samples/cell-exp/3.0.0/pbmc_1k_v3/pbmc_1k_v3_fastqs.tar .`

`tar -xvf pbmc_1k_v3_fastqs.tar`

Now we have:

`
pbmc_1k_v3_fastqs/
    pbmc_1k_v3_S1_L001_I1_001.fastq.gz  
    pbmc_1k_v3_S1_L002_I1_001.fastq.gz
    pbmc_1k_v3_S1_L001_R1_001.fastq.gz  
    pbmc_1k_v3_S1_L002_R1_001.fastq.gz
    pbmc_1k_v3_S1_L001_R2_001.fastq.gz  
    pbmc_1k_v3_S1_L002_R2_001.fastq.gz
`

These `fastq` files are now stored in whatever directory the above command was executed. We can then execute the `cellranger count` function as described on their <a href="https://support.10xgenomics.com/single-cell-gene-expression/software/pipelines/latest/using/count">website</a>.

# %% [markdown]
#### Download data: SmartSeq

For an example of data generated with ***SmartSeq*** technology, we will download this data from the <a href="https://www.ncbi.nlm.nih.gov/geo/">**GEO Database**</a>. We will use a dataset from the seminal paper, ***Single-cell RNA-seq highlights intratumoral heterogeneity in primary glioblastoma*** [(Patel et al., 2015)](https://doi.org/10.1126/science.1254257)

After obtaining the list of accessions from GEO, we will execute the following code to download the data:

# %% [raw]
#!/bin/bash

# create file var
FILE="patel2015_accessions.txt"

all_lines=`cat $FILE`

for line in $all_lines; do fasterq-dump $line; done

# %% [markdown]
### Step 2. Raw read alignment

# %% [markdown]
For the sake of instruction, we will not use CellRanger in this notebook but instead we will start by aligning the raw reads from the ***(Patel et al, 2015) dataset***. in the `fastq` files we just downloaded to the human genome (hg19), which can be downloaded from <a href="https://hgdownload.soe.ucsc.edu/downloads.html">UCSC Genome Browser</a>.

# %% [markdown]
Generate the STAR-compatible reference genome files (only needs to be done once per reference genome)

# %% [raw]
#!/bin/bash

STAR --runThreadN 6 \
--runMode genomeGenerate \
--genomeDir /home/mvinyard/ref/hg19_idx/ \
--genomeFastaFiles /home/mvinyard/ref/fasta/genome.fa \
--sjdbGTFfile /home/mvinyard/ref/genes/genes.gtf \
--sjdbOverhang 99

# %% [markdown]
Perform the alignment

# %% [raw]
#!/bin/bash

ALL_SRR=$(cat /home/mvinyard/patel_2015/patel2015_accessions.txt)
FASTQS=/home/mvinyard/patel_2015/fastqs
OUT_PREFIX=/home/mvinyard/patel_2015/alignment

for CELL in ${ALL_SRR}
do
  echo ${CELL}
  READ_ONE=${FASTQS}/${CELL}_1.fastq
  READ_TWO=${FASTQS}/${CELL}_2.fastq
  STAR --runThreadN 20 \
  --genomeDir /home/mvinyard/ref/hg19/ \
  --readFilesIn ${READ_ONE} ${READ_TWO} \
  --outFileNamePrefix ${OUT_PREFIX}/${CELL} \
  --outSAMtype BAM SortedByCoordinate \
  --outFilterMultimapNmax 2 \
  --outFilterMismatchNmax 3 \
  --alignIntronMax 1 \
  --alignEndsType EndToEnd
done

# %% [markdown]
Other tools not described in detail:

# %% [markdown]
<a href="https://salmon.readthedocs.io/en/latest/index.html">**Salmon**</a> and <a href="https://www.kallistobus.tools/">**kallisto | bustools**</a> can generate feature matrices much faster than the above-used tools. These methods use a pseudoalignment approach. Additionally, <a href="http://bio-bwa.sourceforge.net/">**BWA**</a> can be used for low-divergence sequence alignment. The linked tutorials can be used to process feature matrices for downstream use with the below-mentioned analysis tools. 

# %% [markdown]
### Step 3. Read counting and feature matrix generation

# %% [markdown]
The following bash script was run using <a href="http://www.htslib.org/">samtoools index</a> and <a href="https://htseq.readthedocs.io/en/master/count.html">htseq-count</a> to index and count the reads per feature in each cell.

# %% [raw]
#!/bin/bash

BAMDIR=/home/mvinyard/patel_2015/alignment/
BAMFILES=$(ls ${BAMDIR})
GTF=/home/mvinyard/ref/gencode.v19.annotation.gtf


for FILE in ${BAMFILES}
do
  echo ${FILE}

  _FILE_=${BAMDIR}${FILE}

  samtools index ${_FILE_}
  echo "indexing of alignment complete"
  OUTFILE=$(echo ${FILE} | cut -d'A' -f 1)
  htseq-count ${_FILE_} ${GTF} > /home/mvinyard/patel_2015/counts/${OUTFILE}.count
done

# %% [markdown]
We can also use <a href="http://www.htslib.org/">**samtools index**</a> and <a href="http://bioinf.wehi.edu.au/featureCounts/">**featureCounts**</a>. In general, this second option, **featureCounts** is much faster.

# %% [raw]
#!/bin/bash

BAMDIR=/home/mvinyard/patel_2015/alignment/
BAMFILES=$(ls ${BAMDIR})
GTF=/home/mvinyard/ref/gencode.v19.annotation.gtf

 for FILE in ${BAMFILES}
 do
   echo ${FILE}
   _FILE_=${BAMDIR}${FILE}
   samtools index ${_FILE_}
   echo "indexing of alignment complete"
   OUTFILE=$(echo ${FILE} | cut -d'A' -f 1)
   featureCounts -T 8 -a ${GTF} -o /home/mvinyard/patel_2015/featureCounts/${OUTFILE}.txt ${_FILE_}
 done

# %% [markdown]
Both of these methods generate a text file. Each line contains a feature name and a feature count (and other information, if using featureCounts). We can cut the column containing the counts from each file and paste it together with the genomic coordinates to get a feature matrix, cell x feature. 

# %% [raw]
#!/bin/bash

COUNTDIR=/home/mvinyard/patel_2015/featureCounts/
CELLS=$(ls ${COUNTDIR})
OUTDIR=/home/mvinyard/patel_2015/count_cols

for CELL in ${CELLS}
do  
  cat ${COUNTDIR}${CELL} | cut -f7 > ${OUTDIR}/${CELL}/_count_col.txt
done

cat SRR1294581.txt | cut -f1 > ${OUTDIR}/_coords.txt

paste * > feature_matrix.txt

# %% [markdown]
The output looks something like this (if you only had three cells!):

# %% [raw]
ENSG00000231716.1	0	0	0
ENSG00000234744.1	0	0	0
ENSG00000233156.1	0	0	0
ENSG00000234888.1	0	0	0
ENSG00000215507.6	0	0	0
ENSG00000235014.1	0	0	0
ENSG00000225876.1	0	0	0
ENSG00000251796.1	0	0	0
ENSG00000229238.3	0	0	0
ENSG00000252948.1	0	0	0
ENSG00000233843.1	0	0	0
ENSG00000188399.5	0	0	0
ENSG00000215506.5	0	0	0
ENSG00000224240.1	0	0	0
ENSG00000227629.1	0	0	0
ENSG00000237917.1	0	0	0
ENSG00000231514.1	0	0	0
ENSG00000235857.1	0	0	0
ENSG00000210049.1	0	0	0
ENSG00000211459.2	105	571	3300
ENSG00000210077.1	2	0	0
ENSG00000210082.2	649	3507	2718
ENSG00000209082.1	7	2	0
ENSG00000198888.2	203	404	82
ENSG00000210100.1	2	3	38
ENSG00000210107.1	2	1	8
ENSG00000210112.1	9	4	18
ENSG00000198763.3	81	212	943
ENSG00000210117.1	2	0	0
ENSG00000210127.1	1	2	13
ENSG00000210135.1	1	8	39
ENSG00000210140.1	0	1	5
ENSG00000210144.1	0	9	46
ENSG00000198804.2	920	4036	5264
ENSG00000210151.2	9	23	38
ENSG00000210154.1	18	2	0
ENSG00000198712.1	856	2257	3029
ENSG00000210156.1	35	11	2
ENSG00000228253.1	115	69	83
ENSG00000198899.2	151	277	660
ENSG00000198938.2	84	438	1750
ENSG00000210164.1	1	5	0
ENSG00000198840.2	6	37	377
ENSG00000210174.1	1	4	0
ENSG00000212907.2	60	251	592
ENSG00000198886.2	598	2980	4738
ENSG00000210176.1	2	3	0
ENSG00000210184.1	1	4	0
ENSG00000210191.1	1	3	0
ENSG00000198786.2	550	707	0
ENSG00000198695.2	298	362	0
ENSG00000210194.1	1	4	0
ENSG00000198727.2	478	391	336
ENSG00000210195.2	0	10	0

# %% [markdown]
### Next steps
Now we are ready to move on to one of the supported analysis pipelines: *Scanpy*, *Seurat*, *PAGA*, and *STREAM*.
```
