## Supplementary IPython Notebooks for "*singlecellVR*: interactive visualization of single-cell data in virtual reality": Supplementary_Notebook_3A.ipynb

```
# %% [markdown]
### Supplementary Notebook 3A
---
### Scanpy analysis performed on 1.1 Million cell mouse brain dataset [(Yao et al., 2020)](https://doi.org/10.1101/2020.03.30.015214)

This dataset is shown in **Figure 3A** of the manuscript.

**Preprint**: [A taxonomy of transcriptomic cell types across the isocortex and hippocampal formation](https://www.biorxiv.org/content/10.1101/2020.03.30.015214v1) 

**Raw Data**: [link](https://portal.brain-map.org/atlases-and-data/rnaseq/mouse-whole-cortex-and-hippocampus-10x) 

# %% [markdown]
---
First, we will run the standard preprocessing analysis from scanpy. The data can be downloaded from the link above and with the `matrix.csv` file anyone should be able to reproduce the following analysis. 

# %% [code]
import warnings
warnings.filterwarnings('ignore')

import numpy as np
import pandas as pd
import scanpy as sc

# %% [code]
results_file = 'mousebrain_scanpy.h5ad'  # the file that will store the analysis results

# %% [code]
adata = sc.read_csv('matrix.csv')

# %% [code]
adata.obs = pd.read_csv('./metadata.csv')
adata.obs = adata.obs.set_index('sample_name', drop=True)
adata.obs.head()

# %% [code]
adata

# %% [code]
adata.var.head()

# %% [code]
x = ['exp_component_name', 'platform_label', 'cluster_color', 'cluster_order', 'cluster_label', 'class_color', 'class_order', 'class_label', 'subclass_color', 'subclass_order', 'subclass_label', 'full_genotype_color', 'full_genotype_id', 'full_genotype_label', 'sex_color', 'sex_id', 'donor_sex_label', 'region_color', 'region_id', 'region_label', 'cell_type_accession_color', 'cell_type_accession_id', 'cell_type_accession_label', 'cell_type_alias_color', 'cell_type_alias_id', 'cell_type_alias_label', 'cell_type_alt_alias_color', 'cell_type_alt_alias_id', 'cell_type_alt_alias_label', 'cell_type_designation_color', 'cell_type_designation_id', 'cell_type_designation_label', 'neighborhood_label', 'neighborhood_id', 'neighborhood_color', 'external_donor_name_color', 'external_donor_name_id', 'external_donor_name_label', 'facs_population_plan_color', 'facs_population_plan_id', 'facs_population_plan_label', 'injection_materials_color', 'injection_materials_id', 'injection_materials_label', 'injection_method_color', 'injection_method_id', 'injection_method_label', 'injection_roi_color', 'injection_roi_id', 'injection_roi_label', 'injection_type_color', 'injection_type_id', 'injection_type_label', 'cortical_layer_label', 'outlier_call', 'outlier_type']

for i in x:
    print(i)

# %% [code]
adata.write(results_file)

# %% [code]
adata.var_names_make_unique()  # this is unnecessary if using `var_names='gene_ids'` in `sc.read_10x_mtx`

# %% [code]
sc.pl.highest_expr_genes(adata, n_top=20, )

# %% [code]
sc.pp.filter_cells(adata, min_genes=200)
sc.pp.filter_genes(adata, min_cells=3)

# %% [code]
adata.var['mt'] = adata.var_names.str.startswith('MT-')  # annotate the group of mitochondrial genes as 'mt'
sc.pp.calculate_qc_metrics(adata, qc_vars=['mt'], percent_top=None, log1p=False, inplace=True)
sc.pl.violin(adata, ['n_genes_by_counts', 'total_counts', 'pct_counts_mt'],
             jitter=0.4, multi_panel=True)

# %% [code]
sc.pl.scatter(adata, x='total_counts', y='pct_counts_mt')
sc.pl.scatter(adata, x='total_counts', y='n_genes_by_counts')
adata = adata[adata.obs.n_genes_by_counts < 2500, :]
adata = adata[adata.obs.pct_counts_mt < 5, :]

# %% [code]
sc.pp.normalize_total(adata, target_sum=1e4)
sc.pp.log1p(adata)

# %% [code]
sc.pp.highly_variable_genes(adata, min_mean=0.0125, max_mean=3, min_disp=0.5)
sc.pl.highly_variable_genes(adata)

# %% [code]
sc.pp.scale(adata, max_value=10)

# %% [markdown]
Now that we've done the relatively "standard" Scanpy preprocessing analysis, we can do a bit of data wrangling and adjustment to get things into proper order for preparation wtih `scvr`.

# %% [code]
sc.tl.pca(adata, svd_solver='arpack')
sc.pl.pca(adata, color='Kdm1a')
sc.pl.pca_variance_ratio(adata, log=True)

# %% [code]
sc.pp.neighbors(adata, n_neighbors=10, n_pcs=40)
sc.tl.umap(adata, n_components=3)

# %% [markdown]
Quickly plot some of my favorite genes just to see how it looks:

# %% [code]
sc.pl.umap(adata, color=['Nsd2'])

# %% [code]
sc.pl.umap(adata, color=['Kdm1a'])

# %% [code]
sc.tl.leiden(adata)
sc.tl.louvain(adata)

# %% [code]
adata.raw = adata

# %% [markdown]
Quick save

# %% [code]
adata.write('./mousebrain_scanpy_raw.h5ad')

# %% [markdown]
Let's pull out the high variance genes for annotations in 3D VR.

# %% [code]
highvar = adata.var.loc[adata.var.highly_variable == True][0:10].index # .sort_values('n_cells', ascending = False)[0:10].index
highvar = adata.var.loc[adata.var.highly_variable == True][0:50].sort_values('n_cells', ascending = False)[0:20].index

# %% [markdown]
Plot UMAPs for high-variance genes

# %% [code]
sc.pl.umap(adata, color = highvar)

# %% [code]
adata.var.loc[adata.var.highly_variable == True]

# %% [code]
highvar

# %% [code]
print(results_file)
adata.write(results_file)

# %% [markdown]
Format `adata.obs` to reflect the metadata file

---

# %% [code]
adata

# %% [code]
adata.raw.var.loc[adata.raw.var.highly_variable == True].sort_values('n_cells', ascending = False)[0:10]

# %% [code]
adata.obs

# %% [code]
meta = df[0:3].set_index('Sample ID').T
meta.head()

# %% [code]
adata.obs.index.name = 'Sample ID'
adata.obs[['Cell picking session', 'Cell picking temperature', 'Sex']] = meta[['Cell picking session', 'Cell picking temperature', 'Sex']]

# %% [code]
adata.write(results_file)

# %% [markdown]
---

# %% [code]
adata = sc.read_h5ad('mousebrain_scanpy.h5ad')

# %% [markdown]
Depending on the desired annotations and genes to be visualized by the user, the following command can be executed to create *singlecellVR*-compatible .JSON object. 

`scvr -f mousebrain_scanpy.h5ad -t scanpy -a ANNOTATIONS [-g GENES] [-o OUTPUT]`

# %% [code]
```
