## Supplementary IPython Notebooks for "*singlecellVR*: interactive visualization of single-cell data in virtual reality": Supplementary_Notebook_3B1.ipynb

```
# %% [markdown]
### Supplementary Notebook 3B-1
---
### PAGA analysis performed on a popular mouse hematopoiesis dataset [(Paul et al., 2015)](https://doi.org/10.1016/j.cell.2015.11.013)

This dataset is shown in **Figure 3B** of the manuscript.

# %% [markdown]
Tracing myeloid and erythroid differentiation for data of [Paul et al. (2015)](http://doi.org/10.1016/j.cell.2015.11.013).

# %% [code]
import numpy as np
import pandas as pd
import matplotlib.pyplot as pl
from matplotlib import rcParams
import scanpy as sc

sc.settings.verbosity = 3  # verbosity: errors (0), warnings (1), info (2), hints (3)
sc.logging.print_versions()
results_file = './paga_result/paga3d_paul15.h5ad'

# %% [code]
sc.settings.set_figure_params(dpi=80, frameon=False)  # low dpi (dots per inch) yields small inline figures

# %% [code]
adata = sc.datasets.paul15()

# %% [code]
adata_raw = sc.datasets.paul15()
sc.pp.log1p(adata_raw)
sc.pp.scale(adata_raw)
adata.raw = adata_raw

# %% [code]
adata

# %% [markdown]
Let us work with a higher precision than the default 'float32' to ensure _exactly_ the same results across different computational platforms.

# %% [code]
adata.X = adata.X.astype('float64')  # this is not required and results will be comparable without it

# %% [markdown]
## Preprocessing and Visualization

# %% [markdown]
Apply a simple [preprocessing recipe](https://scanpy.readthedocs.io/en/latest/api/scanpy.api.pp.recipe_zheng17.html).

# %% [code]
sc.pp.recipe_zheng17(adata)

# %% [code]
sc.tl.pca(adata, svd_solver='arpack')

# %% [code]
sc.pl.pca_variance_ratio(adata, log=True, n_pcs=50)

# %% [code]
sc.pp.neighbors(adata, n_neighbors=4, n_pcs=20)

# %% [code]
sc.tl.louvain(adata, resolution=1.0)

# %% [code]
sc.tl.paga(adata, groups='louvain')
sc.pl.paga(adata, plot=True)  # remove `plot=False` if you want to see the coarse-grained graph

# %% [code]
sc.pl.paga(adata, color=['louvain', 'Hba-a2', 'Elane', 'Irf8'])

# %% [code]
sc.pl.paga(adata, color=['louvain', 'Itga2b', 'Prss34', 'Cma1'])

# %% [markdown]
> **Note: In order to use singlecellVR, users need to use at least 3 components for umap**

# %% [code]
sc.tl.umap(adata,n_components=3)

# %% [code]
sc.pl.umap(adata, color=['louvain'])

# %% [code]
sc.tl.rank_genes_groups(adata, 'louvain', method='t-test')
sc.pl.rank_genes_groups(adata, n_genes=25, sharey=False)

# %% [code]
pd.DataFrame(adata.uns['rank_genes_groups']['names'])

# %% [code]
sc.pl.umap(adata, color=pd.DataFrame(adata.uns['rank_genes_groups']['names']).iloc[1,:].tolist(),
           legend_fontsize='x-small',wspace=0.2)

# %% [code]
adata.write(results_file)

# %% [markdown]
> saved `.h5ad` file can be downloaded [here](https://www.dropbox.com/s/qauvu3j0i0t6a3b/paga3d_paul15.h5ad?dl=0)

# %% [markdown]
Depending on the desired annotations and genes to be visualized by the user, the following command can be executed to create *singlecellVR*-compatible .JSON object. 

`scvr -f file.h5ad -t PAGA -a ANNOTATIONS [-g GENES] [-o OUTPUT]`
```
