## Supplementary IPython Notebooks for "*singlecellVR*: interactive visualization of single-cell data in virtual reality": Supplementary_Notebook_3B2.ipynb

````
# %% [markdown]
### Supplementary Notebook 3B2
---
### STREAM analysis performed on a popular mouse hematopoiesis dataset [(Nestorowa et al., 2016)](https://doi.org/10.1182/blood-2016-05-716480)

This dataset is shown in **Figure 3B** of the manuscript.

# %% [markdown]
Datasets can be downloaded [here](https://www.dropbox.com/sh/2vmlbo7sgf5w26v/AABuNn2MATJ8Tm_mpqPZIcw2a?dl=0)   

Data preparation is illustrated [here](https://nbviewer.jupyter.org/github/pinellolab/STREAM/blob/master/tutorial/prep_nestorowa2016.ipynb) 

Ref: Nestorowa, S. et al. A single-cell resolution map of mouse hematopoietic stem and progenitor cell differentiation. Blood 128, e20-31 (2016).

# %% [code]
import stream as st
st.__version__

# %% [code]
st.set_figure_params(dpi=80,style='white',figsize=[5.4,4.8],
                     rc={'image.cmap': 'viridis'})

# %% [markdown]
#### Read in data

# %% [code]
adata=st.read(file_name='./data_nestorowa2016.tsv.gz',workdir='./stream_result')

# %% [markdown]
> **To load and use 10x Genomics single cell RNA-seq data processed with Cell Ranger:**   
(*The variable index can be reset by choosing a different column in `gene.tsv`*)
```python
adata=st.read(file_name='./filtered_gene_bc_matrices/matrix.mtx',
              file_feature='./filtered_gene_bc_matrices/genes.tsv',
              file_sample='./filtered_gene_bc_matrices/barcodes.tsv',
              file_format='mtx',workdir='./stream_result')    
adata.var.index = adata.var[1].values   
```

> **If the Anndata object is already created, to run STREAM, please simply specify work directory:**
```python
st.set_workdir(adata,'./stream_result')
```

# %% [code]
adata.var_names_make_unique()

# %% [code]
adata

# %% [markdown]
#### Read in metadata

# %% [code]
st.add_metadata(adata,file_name='./metadata.tsv')

# %% [code]
adata.obs.head()

# %% [markdown]
> **Alternatively, the step can be divided into two step:**  
(*if `file_name` is not specified, 'unknown' will be genereated for cell label and random color will be used for each label*)
```python
st.add_cell_labels(adata,file_name='./cell_label.tsv.gz')
st.add_cell_colors(adata,file_name='./cell_label_color.tsv.gz')
```

# %% [markdown]
#### Calculate QC

# %% [code]
st.cal_qc(adata,assay='rna')

# %% [code]
st.plot_qc(adata,jitter=0.3,)

# %% [code]
### histogram plots and log-scale are also supported
st.plot_qc(adata,jitter=0.3,log_scale=[0,1,4,5],hist_plot=[0,1,4,5]) 

# %% [code]
st.filter_cells(adata,min_n_features= 100)
st.filter_features(adata,min_n_cells = 5)

# %% [code]
###Normalize gene expression based on library size  
st.normalize(adata,method='lib_size')
###Logarithmize gene expression 
st.log_transform(adata)
###Remove mitochondrial genes 
st.remove_mt_genes(adata)

# %% [markdown]
#### Feature selection

# %% [markdown]
Please check if the blue curve fits the points well. If not, please adjust the parameter **'loess_frac'** (usually by lowering it) until the blue curve fits well. 

# %% [code]
st.select_variable_genes(adata,loess_frac=0.01,n_genes=2000)

# %% [code]
st.select_top_principal_components(adata,feature='var_genes',first_pc=True,n_pc=40)

# %% [markdown]
#### Dimensional reduction

# %% [code]
st.dimension_reduction(adata,method='se',feature='top_pcs',n_components=4,n_neighbors=15,n_jobs=4)

# %% [markdown]
> **Alternatively, using variable genes as features:**
`st.dimension_reduction(adata,method='se',feature='var_genes',n_neighbors=15, n_components=4)`

# %% [code]
st.plot_dimension_reduction(adata,color=['label','Gata1','n_genes'],
                            show_graph=False,show_text=False)

# %% [code]
st.plot_dimension_reduction(adata,color=['label'],
                            show_graph=False,show_text=False,plotly=True)

# %% [markdown]
#### Visualize cells on 2D plane when `n_components>=3` in `st.dimension_reduction()`

# %% [code]
st.plot_visualization_2D(adata,method='umap',n_neighbors=50,
                         color=['label','Gata1','n_genes'],use_precomputed=False)

# %% [markdown]
#### Trajectory inference

# %% [markdown]
* By default, trajectories will be inferred in **the manifold learnt from the step of dimension reduction `st.dimension_reduction()`**


* But it can be also learned directly on **2D visualization plot from `st.plot_visualization_2D()`** by setting `use_vis=True` in st.seed_elastic_principal_graph(). An example can be found [here](https://nbviewer.jupyter.org/github/pinellolab/STREAM/blob/master/tutorial/1.2.use_vis.ipynb)

# %% [code]
st.seed_elastic_principal_graph(adata,n_clusters=20)

# %% [code]
st.plot_dimension_reduction(adata,color=['label','Gata1','n_genes'],n_components=3,show_graph=True,show_text=False)
st.plot_branches(adata,show_text=True)

# %% [markdown]
**`epg_alpha`, `epg_mu`, `epg_lambda` are the three most influential parameters for learning elastic principal graph.**  
- `epg_alpha`: penalizes spurious branching events. **The larger, the fewer branches the function will learn**. (by default, `epg_alpha=0.02`)
- `epg_mu`: penalizes the deviation from harmonic embedding, where harmonicity assumes that each node is the mean of its neighbor nodes. **The larger, the more edges the function will use to fit into points(cells)** (by default, `epg_mu=0.1`) 
- `epg_lambda`: penalizes the total length of edges. **The larger, the 'shorter' curves the function will use to fit into points(cells)** and the fewer points(cells) the curves will reach. (by default, `epg_lambda=0.02`)

> **'epg_trimmingradius'** can help get rid of noisy points (by defalut `epg_trimmingradius=Inf`)   
e.g. `st.elastic_principal_graph(adata,epg_trimmingradius=0.1)`  

# %% [code]
st.elastic_principal_graph(adata,epg_alpha=0.02,epg_mu=0.05,epg_lambda=0.01)

# %% [code]
st.plot_dimension_reduction(adata,color=['label','Gata1','n_genes'],n_components=3,show_graph=True,show_text=False)
st.plot_branches(adata,show_text=False)

# %% [markdown]
#### Adjusting trajectories (optional)

# %% [markdown]
* Finetune branching event:  
```python
st.optimize_branching(adata,incr_n_nodes=30)
st.plot_dimension_reduction(adata,show_graph=True,show_text=False)
st.plot_branches(adata,show_text=False)
```
* Prune trivial branches:  
```python
st.prune_elastic_principal_graph(adata,epg_collapse_mode='EdgesNumber',epg_collapse_par=2)
st.plot_dimension_reduction(adata,show_graph=True,show_text=False)
st.plot_branches(adata,show_text=False)
```

* Shift branching node:  
```python
st.shift_branching(adata,epg_shift_mode='NodeDensity',epg_shift_radius=0.1,epg_shift_max=3)
st.plot_dimension_reduction(adata,show_graph=True,show_text=False)
st.plot_branches(adata,show_text=False)
```

# %% [code]
###Extend leaf branch to reach further cells 
st.extend_elastic_principal_graph(adata, epg_ext_mode='WeigthedCentroid',epg_ext_par=0.8)
st.plot_dimension_reduction(adata,color=['label'],n_components=3,show_graph=True,show_text=True)
st.plot_branches(adata,show_text=True)

# %% [code]
st.plot_dimension_reduction(adata,color=['label'],n_components=3,show_graph=True,show_text=True,plotly=True)

# %% [code]
st.plot_visualization_2D(adata,n_neighbors=50,color=['label','Gata1','branch_id_alias','S1_pseudotime'],fig_ncol=4)

# %% [code]


# %% [markdown]
#### Trajectory visualization

# %% [markdown]
##### flat tree

# %% [code]
st.plot_flat_tree(adata,color=['label','branch_id_alias','S3_pseudotime'],
                  dist_scale=0.5,show_graph=True,show_text=True)

# %% [markdown]
##### stream plot at single cell level

# %% [code]
st.plot_stream_sc(adata,root='S1',color=['label','Gata1'],
                  dist_scale=0.3,show_graph=True,show_text=True)

# %% [markdown]
##### stream plots

# %% [code]
st.plot_stream(adata,root='S1',color=['label','Gata1'])

# %% [markdown]
**Some useful parameters to finetune the appearance of stream plots:**
* `dist_scale`: controls the width of STREAM plot branches.
* `factor_num_win`: controls the resolution of STREAM plot.
* `preference`: controls the order of branches
* `log_scale`: shows stream plot in log scale to zoom in on thin branches

# %% [code]
st.plot_stream(adata,root='S1',color=['label','Gata1'],factor_min_win=1.5)

# %% [markdown]
#### Marker genes detection

# %% [markdown]
`marker_list` defines the list of genes to scan. If not specified, by default it uses all available genes. It might be time-consuming.  

Here we only include variable genes.

# %% [markdown]
##### 1) detect marker genes for each leaf branch

# %% [code]
st.detect_leaf_markers(adata,marker_list=adata.uns['var_genes'],cutoff_zscore=1.0,cutoff_pvalue=0.01,
                       root='S1',n_jobs=4)

# %% [code]
adata.uns['leaf_markers_all'].head()

# %% [code]
adata.uns['leaf_markers'][('S3','S5')].head()

# %% [markdown]
##### 2) detect transition genes for each branch

# %% [code]
st.detect_transition_markers(adata,marker_list=adata.uns['var_genes'],cutoff_spearman=0.4,cutoff_logfc=0.25,
                             root='S1',n_jobs=4)

# %% [code]
adata.uns['transition_markers'][('S3','S4')].head()

# %% [code]
st.plot_transition_markers(adata,fig_size=(10,5))

# %% [markdown]
##### 3) detect marker genes that are differentially expressed between pairs of branches

# %% [code]
st.detect_de_markers(adata,marker_list=adata.uns['var_genes'],cutoff_zscore=1,cutoff_logfc=0.25,
                     root='S1',n_jobs=4)

# %% [code]
adata.uns['de_markers_greater'][(('S3', 'S4'), ('S3', 'S5'))].head()

# %% [code]
adata.uns['de_markers_less'][(('S3', 'S4'), ('S3', 'S5'))].head()

# %% [code]
st.plot_de_markers(adata)

# %% [markdown]
##### 4) detect cell population-specific markers

```python
st.detect_markers(adata,ident='label',marker_list=adata.uns['var_genes'],cutoff_zscore=1.0,cutoff_pvalue=0.01)
```

# %% [markdown]
#### Save results

# %% [code]
st.write(adata,file_name='stream_result.pkl')

# %% [markdown]
To read back the saved `.pkl` file
```python
adata = st.read('./stream_result/stream_result.pkl')
```

# %% [markdown]
##### Additionally, STREAM analysis result can be also saved
* for single cell VR website (http://singlecellvr.com/)
```python
st.save_vr_report(adata,ann_list=['label','S1_pseudotime'],gene_list=['Gata1','Epx','Car2'])
```
* for STREAM website (http://stream.pinellolab.org/)
```python
st.save_web_report(adata,n_genes=5,
                    title='Nestorowa, S. et al. 2016',
                    description='This scRNA-seq dataset contains 1656 cells and 4768 genes from mouse hematopoietic stem and progenitor cell differentiation. A single-cell resolution map of mouse hematopoietic stem and progenitor cell differentiation. Blood 128, e20-31 (2016).',
                    starting_node='S1',command_used='1.2.STREAM_scRNA-seq (Multifurcation)')
```

# %% [markdown]
Depending on the desired annotations and genes to be visualized by the user, the following command can be executed to create *singlecellVR*-compatible .JSON object. 

`scvr -f stream_result.pkl -t STREAM -a ANNOTATIONS [-g GENES] [-o OUTPUT]`
````
