## Supplementary IPython Notebooks for "*singlecellVR*: interactive visualization of single-cell data in virtual reality": Supplementary_Notebook_3C1.ipynb

````
# %% [markdown]
### Supplementary Notebook 3C-1
---
### Episcanpy [(Danese et al., 2019)](https://doi.org/10.1101/648097) analysis performed on the 10x PBMC scATAC-seq dataset 

The processed result of this datasest is shown in **Figure 3C (left)** of the manuscript.

**Preprint**: [EpiScanpy: integrated single-cell epigenomic analysis](https://www.biorxiv.org/content/10.1101/648097v1) 

**Raw Data**: [link](https://support.10xgenomics.com/single-cell-atac/datasets/1.2.0/atac_pbmc_10k_nextgem) 

# %% [markdown]
```
wget https://cf.10xgenomics.com/samples/cell-atac/1.2.0/atac_pbmc_10k_nextgem/atac_pbmc_10k_nextgem_filtered_peak_bc_matrix.tar.gz
```

```
tar xzvf atac_pbmc_10k_nextgem_filtered_peak_bc_matrix.tar.gz
cd atac_pbmc_10k_nextgem_filtered_peak_bc_matrix
gzip *
``` 

# %% [code]
wd = "/home/mvinyard/scvr_polished_nbs/data/fig3c/filtered_peak_bc_matrix/"


matrix     = wd + 'matrix.mtx.gz'
cell_names = wd + 'barcodes.tsv.gz'
var_names  = wd + 'features.tsv.gz'

# %% [code]
import warnings
warnings.filterwarnings('ignore')

import episcanpy as epi
import pandas as pd
import numpy as np
import scanpy as sc

# %% [code]
adata = epi.read_mtx(matrix).T

# %% [code]
adata.obs = pd.read_csv(cell_names, header = None, sep='\t').rename({0:'barcodes'}, axis=1)
adata.var = pd.read_csv(var_names, header = None, sep='\t').rename({0:'chr',1:'start',2:'stop'}, axis=1)
adata

# %% [code]
epi.pp.commonness_features(adata)

# %% [code]
epi.pp.coverage_cells(adata)

# %% [code]
adatabin = epi.pp.binarize(adata, copy=True)

# %% [code]
epi.pp.coverage_cells(adatabin)

# %% [code]
sc.pp.pca(adata)
sc.pp.neighbors(adata)

# %% [code]
sc.tl.pca(adata)
sc.tl.tsne(adata)
sc.tl.louvain(adata)

# %% [code]
sc.tl.umap(adata, n_components=3)

# %% [code]
sc.pl.pca(adata, color='louvain')
sc.pl.tsne(adata, color='louvain')
sc.pl.umap(adata, color='louvain')

# %% [code]
adata

# %% [code]
results_file = './10x_10kcells_pbmc.h5ad'
adata.write(results_file)

# %% [markdown]
Depending on the desired annotations and genes to be visualized by the user, the following command can be executed to create *singlecellVR*-compatible .JSON object. 

`scvr -f ./10x_10kcells_pbmc.h5ad -t scanpy -a ANNOTATIONS [-g GENES] [-o OUTPUT]`
````
