## Supplementary IPython Notebooks for "*singlecellVR*: interactive visualization of single-cell data in virtual reality": Supplementary_Notebook_3C2.ipynb

````
# %% [markdown]
### Supplementary Notebook 3C-2
---
### STREAM analysis performed on a human hematopoeitic differentiation dataset [(Buenrostro et al., 2018)](https://doi.org/10.1016/j.cell.2018.03.074)

The processed result of this datasest is shown in **Figure 3C (right)** of the manuscript.

**Publication**: [Integrated Single-Cell Analysis Maps the Continuous Regulatory Landscape of Human Hematopoietic Differentiation](https://www.cell.com/cell/fulltext/S0092-8674(18)30446-X?_returnURL=https%3A%2F%2Flinkinghub.elsevier.com%2Fretrieve%2Fpii%2FS009286741830446X%3Fshowall%3Dtrue) 

**Raw Data**: [link](https://www.dropbox.com/sh/xekjl6x2yw4eizf/AADkPqRyXFcT5XrQJZiE5hO9a?dl=0)

**The data preparation**: [link](https://nbviewer.jupyter.org/github/pinellolab/STREAM/blob/master/tutorial/prep_buenrostro_2018.ipynb) 

# %% [code]
import stream as st
st.__version__

# %% [code]
st.set_figure_params(dpi=80,style='white',figsize=[5.4,4.8],
                     rc={'image.cmap': 'viridis'})

# %% [code]
adata=st.read(file_name='./mat.tsv.gz',workdir='./result_atac')

# %% [code]
adata

# %% [markdown]
> **To load and use 10x Genomics single cell ATAC-seq data processed with Cell Ranger:**   
```python
adata=st.read(file_name='./filtered_peak_bc_matrix/matrix.mtx',
              file_sample='./filtered_peak_bc_matrix/barcodes.tsv',
              file_feature='./filtered_peak_bc_matrix/peaks.bed',
              file_format='mtx',workdir='./result_atac') 
adata.var.index = adata.var[0].astype(str) + '_' + adata.var[1].astype(str) +'_' + adata.var[2].astype(str)
```

# %% [code]
st.add_cell_labels(adata,file_name='./cell_label.tsv.gz')
st.add_cell_colors(adata,file_name='./cell_label_color.tsv.gz')

# %% [code]
st.cal_qc(adata,assay='atac')

# %% [code]
st.plot_qc(adata,jitter=0.3)

# %% [code]
st.plot_qc(adata,log_scale=[0,1,3,4],hist_plot=range(6),jitter=0.3)

# %% [code]
st.filter_cells(adata,min_n_features=100)
st.filter_features(adata,min_n_cells=20)

# %% [markdown]
#### LSI transformation

# %% [code]
adata.X[adata.X>0] = 1 ###binarization
st.normalize(adata,method='tf_idf')

# %% [code]
adata

# %% [markdown]
#### Feature selection

# %% [code]
st.select_top_principal_components(adata,first_pc=False,n_pc=50)

# %% [markdown]
#### Dimension reduction

# %% [code]
st.dimension_reduction(adata,method='se',feature='top_pcs',n_components=5,n_neighbors=100,n_jobs=4)

# %% [code]
st.plot_dimension_reduction(adata,color=['label'],show_graph=False,show_text=False,)

# %% [code]
st.plot_visualization_2D(adata,n_neighbors=100,color=['label'],use_precomputed=False)

# %% [markdown]
#### Trajectory inference

# %% [markdown]
* By default, trajectories will be inferred in **the manifold learnt from the step of dimension reduction `st.dimension_reduction()`**


* But it can be also learned directly on **2D visualization plot from `st.plot_visualization_2D()`** by setting `use_vis=True` in st.seed_elastic_principal_graph(). An example can be found [here](https://nbviewer.jupyter.org/github/pinellolab/STREAM/blob/master/tutorial/1.2.use_vis.ipynb)

# %% [code]
st.seed_elastic_principal_graph(adata,n_clusters=10)

# %% [code]
st.plot_dimension_reduction(adata,color=['label'],show_graph=True,show_text=False)
st.plot_branches(adata,show_text=True)

# %% [code]


# %% [markdown]
**`epg_alpha`, `epg_mu`, `epg_lambda` are the three most influential parameters for learning elastic principal graph.**  
- `epg_alpha`: penalizes spurious branching events. **The larger, the fewer branches the function will learn**. (by default, `epg_alpha=0.02`)
- `epg_mu`: penalizes the deviation from harmonic embedding, where harmonicity assumes that each node is the mean of its neighbor nodes. **The larger, the more edges the function will use to fit into points(cells)** (by default, `epg_mu=0.1`) 
- `epg_lambda`: penalizes the total length of edges. **The larger, the 'shorter' curves the function will use to fit into points(cells)** and the fewer points(cells) the curves will reach. (by default, `epg_lambda=0.02`)

> **'epg_trimmingradius'** can help get rid of noisy points (by defalut `epg_trimmingradius=Inf`)   
e.g. `st.elastic_principal_graph(adata,epg_trimmingradius=0.1)`  

# %% [code]
st.elastic_principal_graph(adata,epg_alpha=0.01,epg_mu=0.05,epg_lambda=0.01)

# %% [code]
st.plot_visualization_2D(adata,n_neighbors=30,color=['label','branch_id_alias'],fig_ncol=4)

# %% [code]
st.plot_dimension_reduction(adata,color=['label'],show_graph=True,show_text=False)
st.plot_branches(adata,show_text=True)

# %% [code]
###Extend leaf branch to reach further cells 
st.extend_elastic_principal_graph(adata, epg_ext_mode='WeigthedCentroid',epg_ext_par=0.8)
st.plot_dimension_reduction(adata,color=['label'],show_graph=True,show_text=True)
st.plot_branches(adata,show_text=True)

# %% [markdown]
#### Trajectory visualization

# %% [code]
st.plot_flat_tree(adata,color=['label','branch_id_alias','S2_pseudotime'],
                  dist_scale=0.5,show_graph=False,show_text=False)

# %% [code]
st.plot_flat_tree(adata,color=['label','branch_id_alias','S2_pseudotime'],
                  dist_scale=0.5,show_graph=True,show_text=True)

# %% [code]
st.plot_stream_sc(adata,root='S2',color=['label'],
                  dist_scale=0.5,show_graph=True,show_text=True)

# %% [code]
st.plot_stream(adata,root='S2',color=['label'])

# %% [code]
st.plot_stream(adata,root='S2',color=['label'],dist_scale=1.5)

# %% [markdown]
#### Marker peaks detection

# %% [code]
st.detect_leaf_markers(adata,marker_list=adata.var[adata.var['pct_cells']>0.1].index[:1000],
                       cutoff_zscore=1.0,cutoff_pvalue=0.01,
                       root='S1',n_jobs=4)

# %% [code]
adata.uns['leaf_markers_all'].head()

# %% [code]
st.plot_stream_sc(adata,root='S2',color=['label','chr1_153316244_153316744','chr1_22192536_22193036','chr1_59221069_59221569'],
                  dist_scale=0.5,show_graph=True,show_text=True)

# %% [code]


# %% [code]
st.plot_stream(adata,root='S2',color=['label','chr1_153316244_153316744','chr1_22192536_22193036','chr1_59221069_59221569'],
               dist_scale=1.5)

# %% [markdown]
#### Save results

# %% [code]
st.write(adata,file_name='stream_atac_peaks.pkl')

# %% [markdown]
Depending on the desired annotations and genes to be visualized by the user, the following command can be executed to create *singlecellVR*-compatible .JSON object. 

`scvr -f ./stream_atac_peaks.pkl -t STREAM -a ANNOTATIONS [-g GENES] [-o OUTPUT]`
````
