## Supplementary IPython Notebooks for "*singlecellVR*: interactive visualization of single-cell data in virtual reality": Supplementary_Notebook_3D.ipynb

```
# %% [markdown]
### Supplementary Notebook 3D
---
### Seurat [(Stuart et al., 2019)](https://doi.org/10.1016/j.cell.2019.05.031) analysis performed on the 10x PBMC scATAC-seq dataset 

The processed result of this datasest is shown in **Figure 3D** of the manuscript.

**Publication**: [Comprehensive Integration of Single-Cell Data](https://www.cell.com/cell/fulltext/S0092-8674(19)30559-8) 

**Data**: [link](https://support.10xgenomics.com/single-cell-multiome-atac-gex/datasets/1.0.0/pbmc_granulocyte_sorted_10k)

# %% [code]
options(warn=-1)
library(Seurat)
library(ggplot2)
library(patchwork)

# %% [code]
peaks <- Read10X_h5('atac_v1_pbmc_10k_filtered_peak_bc_matrix.h5')

# %% [code]
activity.matrix <- CreateGeneActivityMatrix(peak.matrix = peaks, 
    annotation.file = '/data/pinello/COMMON_DATA/Homo_sapiens.GRCh37.87.gtf', 
    seq.levels = c(1:22, "X", "Y"), upstream = 2000, verbose = TRUE)

# %% [code]
pbmc.atac <- CreateSeuratObject(counts = peaks, assay = "ATAC", project = "10x_ATAC")
pbmc.atac[["ACTIVITY"]] <- CreateAssayObject(counts = activity.matrix)
meta <- read.table("atac_v1_pbmc_10k_singlecell.csv", sep = ",", header = TRUE, row.names = 1, 
    stringsAsFactors = FALSE)
meta <- meta[colnames(pbmc.atac), ]
pbmc.atac <- AddMetaData(pbmc.atac, metadata = meta)
pbmc.atac <- subset(pbmc.atac, subset = nCount_ATAC > 5000)
pbmc.atac$tech <- "atac"

# %% [code]
DefaultAssay(pbmc.atac) <- "ACTIVITY"
pbmc.atac <- FindVariableFeatures(pbmc.atac)
pbmc.atac <- NormalizeData(pbmc.atac)
pbmc.atac <- ScaleData(pbmc.atac)

# %% [code]
DefaultAssay(pbmc.atac) <- "ATAC"
VariableFeatures(pbmc.atac) <- names(which(Matrix::rowSums(pbmc.atac) > 100))
pbmc.atac <- RunLSI(pbmc.atac, n = 50, scale.max = NULL)
pbmc.atac <- RunUMAP(pbmc.atac, reduction = "lsi", dims = 1:50, n.components = 3)

# %% [code]
pbmc.rna <- readRDS("./pbmc_10k_v3.rds")
pbmc.rna$tech <- "rna"

# %% [code]
p1 <- DimPlot(pbmc.atac, reduction = "umap") + NoLegend() + ggtitle("scATAC-seq")
p2 <- DimPlot(pbmc.rna, group.by = "celltype", label = TRUE, repel = TRUE) + NoLegend() + ggtitle("scRNA-seq")
p1 + p2

# %% [code]
transfer.anchors <- FindTransferAnchors(reference = pbmc.rna, query = pbmc.atac, features = VariableFeatures(object = pbmc.rna), 
    reference.assay = "RNA", query.assay = "ACTIVITY", reduction = "cca")

# %% [code]
celltype.predictions <- TransferData(anchorset = transfer.anchors, refdata = pbmc.rna$celltype, 
    weight.reduction = pbmc.atac[["lsi"]])
pbmc.atac <- AddMetaData(pbmc.atac, metadata = celltype.predictions)

# %% [code]
pbmc.atac.filtered <- subset(pbmc.atac, subset = prediction.score.max > 0.5)
pbmc.atac.filtered$predicted.id <- factor(pbmc.atac.filtered$predicted.id, levels = levels(pbmc.rna))  # to make the colors match
p1 <- DimPlot(pbmc.atac.filtered, group.by = "predicted.id", label = TRUE, repel = TRUE) + ggtitle("scATAC-seq cells") + 
    NoLegend() + scale_colour_hue(drop = FALSE)
p2 <- DimPlot(pbmc.rna, group.by = "celltype", label = TRUE, repel = TRUE) + ggtitle("scRNA-seq cells") + 
    NoLegend()
p1 + p2

# %% [code]
# note that we restrict the imputation to variable genes from scRNA-seq, but could impute the
# full transcriptome if we wanted to
genes.use <- VariableFeatures(pbmc.rna)
refdata <- GetAssayData(pbmc.rna, assay = "RNA", slot = "data")[genes.use, ]

# refdata (input) contains a scRNA-seq expression matrix for the scRNA-seq cells.  imputation
# (output) will contain an imputed scRNA-seq matrix for each of the ATAC cells
imputation <- TransferData(anchorset = transfer.anchors, refdata = refdata, weight.reduction = pbmc.atac[["lsi"]])

# this line adds the imputed data matrix to the pbmc.atac object
pbmc.atac[["RNA"]] <- imputation
coembed <- merge(x = pbmc.rna, y = pbmc.atac)

# Finally, we run PCA and UMAP on this combined object, to visualize the co-embedding of both
# datasets
coembed <- ScaleData(coembed, features = genes.use, do.scale = FALSE)
coembed <- RunPCA(coembed, features = genes.use, verbose = FALSE)
coembed <- RunUMAP(coembed, dims = 1:30, n.components = 3)
coembed$celltype <- ifelse(!is.na(coembed$celltype), coembed$celltype, coembed$predicted.id)

# %% [code]
p1 <- DimPlot(coembed, group.by = "tech")
p2 <- DimPlot(coembed, group.by = "celltype", label = TRUE, repel = TRUE)
p1 + p2

# %% [code]
devtools::install_github(repo = "mojaveazure/loomR", ref = "develop")

# %% [code]
install.packages("reticulate")

# %% [code]
library(reticulate)

# %% [code]
as.loom

# %% [code]
pbmc_ad <- Convert(from = coembed, to = "anndata", filename = "seurat_10xpbmc_coembed.h5ad")

# %% [code]
coembed.loom <- as.loom(coembed, filename = "./seurat_10xpbmc_coembed2.loom", verbose = FALSE)

# %% [code]
coembed.loom$close_all()

# %% [code]
saveRDS(coembed, "./pbmc_10k_v3_coemb.rds")

# %% [code]
pbmc.rna

# %% [code]
pbmc.atac.filtered

# %% [code]
pbmc.rna$observed

# %% [code]
genes.use

# %% [code]
pbmc.rna$

# %% [code]
coembed$

# %% [code]
coembed@graphs <- list()

# %% [code]
coembed = FindVariableFeatures(coembed)

# %% [code]
coembed$orig.ident = NULL

# %% [code]
coembed.loom <- as.loom(coembed, filename = "./seurat_10xpbmc_coembed.loom", verbose = TRUE)

# %% [code]
coembed.loom$close_all()

# %% [code]
coembed@meta.data

# %% [code]
for(j in 1:ncol(coembed@meta.data)){
	if(is.factor(coembed@meta.data[,j]) == T){
	coembed@meta.data[,j] = as.character(coembed@meta.data[,j]) # Force the variable type to be character
	coembed@meta.data[,j][is.na(coembed@meta.data[,j])] <- "N.A"
}
if(is.character(coembed@meta.data[,j]) == T){
	coembed@meta.data[,j][is.na(coembed@meta.data[,j])] <- "N.A"
 }
}

# %% [code]
coembed$

# %% [code]
coembed$orig.ident = NONE

# %% [markdown]
Depending on the desired annotations and genes to be visualized by the user, the following command can be executed to create *singlecellVR*-compatible .JSON object. 

`scvr -f ./seurat_10xpbmc_coembed.loom -t seurat -a ANNOTATIONS [-g GENES] [-o OUTPUT]`
```
