## Supplementary IPython Notebooks for "*singlecellVR*: interactive visualization of single-cell data in virtual reality": Supplementary_Notebook_4.ipynb

```
# %% [markdown]
### Supplementary Notebook 4
---
### SingleCellVR Tutorial : RNA Velocity Preprocessing

This is a complete tutorial for processing raw scRNA-seq data into a velocity map ready for upload onto [SingleCellVR.com](https://singlecellvr.herokuapp.com/).
Due to the amount of raw data collection/processing within the early stages of velocity analysis, users will need access to a remote server that allocates memory and storage for their respective dataset. 

### The main modules/packages used for this analysis are:
1. [velocyto](http://velocyto.org/)
2. [STAR](https://physiology.med.cornell.edu/faculty/skrabanek/lab/angsd/lecture_notes/STARmanual.pdf)
3. [scVelo](https://scvelo.readthedocs.io/)

**NOTE**: There are potential difficulties when downloading the velocyto package- please refer to the [velocyto team github](https://github.com/velocyto-team/velocyto.py).

# %% [markdown]
### Velocyto analysis can only be done on these particular samples:
1. Smartseq
2. Dropseq and/or inDrops
3. Chromium

# %% [markdown]
### Samples can be downloaded from [GEO](https://www.ncbi.nlm.nih.gov/geo/), where users can also receive specifications about the size of files and types of samples there are within a dataset.
[Here](https://www.ncbi.nlm.nih.gov/Traces/study/?acc=PRJNA322317&o=acc_s%3Aa) are samples/accession information from GEO for the dataset used within [Nestorowa et. al, 2016](https://ashpublications.org/blood/article/128/8/e20/35749/A-single-cell-resolution-map-of-mouse).

# %% [markdown]


# %% [markdown]
This [barcodes.txt](https://drive.google.com/file/d/1hgKwaF6Ib1tHgy1wjzxW8-gNyGPSCsJP/view?usp=sharing) includes the SRR number, the cell types, and the cluster group for the Nestorowa dataset. This file will help with coloring and labeling for cells when it is time for visualizing the data. 

# %% [markdown]


# %% [markdown]
The following shell scripts can take multiple hours/days to run, so a command for uploading scripts to the background of your remote server is necessary: 

# %% [code]
nohup mycommand..... > fileNameProgress.out 2>&1 &
#.out can be opened so you can manually view the progress of your command periodically. 

# %% [markdown]


# %% [markdown]


# %% [markdown]
In the terminal, all shell scripts can be run using: 

# %% [code]
sh file.sh

# %% [markdown]


# %% [markdown]


# %% [markdown]
### FastqDump.sh

# %% [code]
!/bin/bash #make sure to run this script in whichever directory you want your samples to be downloaded into. 

BSUB -q bigmulti BSUB -core 5 BSUB -M 64000 BSUB -R rusage[mem=64000] #specs for the particular VPN I used

sample_ID="/filepath/to/cellSampleNames.txt" #this file shoould just be a list of all GEO SRR samples within nestorowaBarcodes.txt

for sample in $(cat $sample_ID); do
  echo fastq-dump --gzip $sample;
        fastq-dump --gzip ${sample} #gzip step is added to make all samples downloaded as file.fastq.gz
done 

# %% [markdown]
Fastq-dump can take approximately one week depending on how many RNA-seq files need to be retrieved for a dataset. 

# %% [markdown]


# %% [markdown]
### fastqToSam.sh
Script edited from this [scRNA-seq pipeline](https://github.com/JunyueC/sci-CAR_analysis/blob/master/scRNA_seq_pipe/scRNA_seq_pipeline.sh), but edited to apply to single-end sequencing data (Nestorowa was using single-end fastq files). 

**NOTE**: Barcodes are extremeley necessary if you want to include coloring/clustering for your velocity file.

# %% [code]
#make sure to load/import samtools for the remaining shell scripts

!/bin/bash 

#BSUB -q bigmulti #BSUB -core 5 #BSUB -M 64000 #BSUB -R rusage[mem=64000]

fastq_folder="/filepath/to/fastq/"
input_folder=$fastq_folder

all_output_folder="/filepath/to/output/"

sample_ID="/filepath/to/sampleNames.txt" 

gtf_file="/filepath/to/genome_file.gtf" #this can be found on ensembl.org and can be obtained using !wget command

core=8 #memory specs 

cutoff=200 #see STAR module for information on this amount

index="/filepath/to/genomeIndex/" #location for genome index which can be retrieved through STAR

script_folder="/filepath/to/scriptFile/" #change back to the script path if needed
script_path=/filepath/to/scriptfile

python_path="/filepath/to/anaconda3/bin"

module load STAR/2.5.4b-foss-2016b #change the specs after STAR/ depending on the version you import

STAR_output_folder=$all_output_folder/STAR_alignment
filtered_sam_folder=$all_output_folder/filtered_sam
rmdup_sam_folder=$all_output_folder/rmdup_sam

mkdir -p $STAR_output_folder
#remove the index from the memory
#STAR --genomeDir $index --genomeLoad Remove
#start the alignment
echo $input_folder
for sample in $(cat $sample_ID); do
  echo Aligning $sample;
  STAR --runThreadN $core --outSAMstrandField intronMotif --genomeDir $index \
  --readFilesCommand zcat \
  --readFilesIn $input_folder/${sample}.fastq --outFileNamePrefix \
  $STAR_output_folder/${sample} #--genomeLoad LoadAndKeep;
done 
#remember to include the readFilesCommand to gunzip the files
STAR --genomeDir $index --genomeLoad Remove
echo "All alignment done."

# %% [markdown]
The fastqToSam step should take about 1-2 days. Once complete you should have fastq files in a human-readable format (sam). These need to be converted to binary-format (bam) and aligned to proceed with velocyto analysis.

# %% [markdown]


# %% [markdown]
### samToBam.sh

# %% [code]
!/bin/bash

#BSUB -q long

ls *.sam | xargs -n 1 -I {} sh -c 'samtools view -F 4 -b {} > {}.bam' 

# %% [markdown]


# %% [markdown]
### bamToSortedBam.sh

# %% [code]
!/bin/bash

#BSUB -q long #BSUB -n 4 #BSUB -M 7000

 for f in *bam
  do samtools sort -@ 4 -o ${f/.bam/sorted.bam} $f
 done

# %% [markdown]


# %% [markdown]
### sortedBamRename.sh
Filenames will look long and messy at this point, so this renaming step will clean everything up.

# %% [code]
!/bin/bash

#BSUB -q long

for f in *.out.samsorted.bam; do
    mv -- "$f" "${f%.out.samsorted.bam}sorted.bam"
done

# %% [markdown]


# %% [markdown]
### SRR to CellID rename

# %% [markdown]
First, convert excel sheet to dictionary and save. This can be done in a jupyter notebook. 

# %% [code]
import pandas as pd
import json

df = pd.read_csv("/Users/Desktop/rdc/SRRCellID.txt") #column 1: SRR names, column 2: cell names
dictDf = df.to_dict() #converts SRR names to keys and cell names to values (dictionary)


# Serialize data into file:
json.dump( dictDf, open( "dictSRRCellID.json", 'w' ) )

# %% [markdown]
Next, this is a shell script that will iterate through the dictionary, finding matching file names and changing them to the cell names. 

# %% [code]
!/bin/bash

#BSUB -q long

import os
import json

os.chdir('/File/path/to/all/.sorted.bam)
         
# Read data from file:
names = json.load( open( "/File/path/to/dictSRRCellID.json" ) )

i = 0
root = os.getcwd()
for filename in os.listdir('.'):
         name = names[filename]
except KeyError: 
    # Filename was not found. Skip. This shouldn't happen, but will prevent your script from breaking for whatever reason. 
    print("Filename "+filename+" not found")
    continue    

# %% [markdown]
### Velocyto step: sortedBamToLoom.sh
Have this script located where the sorted bam files are. 

# %% [code]
!/bin/bash
#BSUB -q bigmulti #BSUB -core 5 #BSUB -M 64000 #BSUB -R rusage[mem=64000]
velocyto run-smartseq2 -o /filepath/to/ouputFolder/ -e *.bam genome_file.gtf #same gtf used in fastqToSam.sh step

# %% [markdown]
Within the outputFolder, there should be your velocity.loom file, which contains the spliced and unspliced counts for your scRNA-seq data.

# %% [markdown]


# %% [markdown]


# %% [markdown]
In a jupyter notebook, convert the cell cluster names within your nestorowaBarcodes.txt file to a numpy.ndarray and add it as a key to your .loom file. Then you will be able to apply coloring and labelling to the file using the scVelo tutorial below. 

# %% [markdown]


# %% [markdown]
### The 'absolute_velocity_umap' key is necessary for uploading the velocity file successfully to SingleCellVR.com.  [Here](https://github.com/qinqian/singlecellvr/blob/master/examples/velocity_3d_scale.ipynb) is a notebook with steps for creating the key. 

# %% [markdown]


# %% [markdown]
### To further prepare the loom file for uploading onto SingleCellVR.com, follow the scVelo tutorial [here](https://github.com/pinellolab/singlecellvr/blob/master/examples/velocity_3d.ipynb).
```
