## Supplementary IPython Notebooks for "*singlecellVR*: interactive visualization of single-cell data in virtual reality": Supplementary_Notebook_S3A.ipynb

```
# %% [markdown]
### Supplementary Notebook S1A
---
### Single-cell transcriptomics of 20 mouse organs creates a *Tabula Muris* [(The Tabula Muris Consortium., Overall coordination., Schaum, N. et al., 2020)](https://doi.org/10.1101/665307)

The processed result of this datasest is shown in **Supplementary Figure 1A** of the manuscript.

**Public**: [Single-cell transcriptomics of 20 mouse organs creates a *Tabula Muris*](https://www.nature.com/articles/s41586-018-0590-4) 

**Raw Data**: [link](http://tabula-muris.ds.czbiohub.org/) 

# %% [code]
import anndata as a
import pandas as pd
import scanpy as sc
import matplotlib as mpl
import matplotlib.pyplot as plt
from matplotlib import cm
from mpl_toolkits.mplot3d import Axes3D

# %% [code]
adata = a.read_h5ad('/data/pinello/PEOPLE/michael/tabula-muris.h5ad')
adata

# %% [code]
sc.pp.highly_variable_genes(adata)

# %% [code]
genes = adata.var.loc[adata.var['highly_variable'] == True].sort_values('dispersions_norm', ascending=False)[0:100].reset_index()['index']
genes.to_csv('tabula_muris-genes.txt',sep='\n', index=False, header=False)
genes

# %% [code]
adata

# %% [code]
sc.pp.neighbors(adata, n_neighbors=4, n_pcs=20)
sc.tl.umap(adata, n_components=3)

# %% [code]
adata.obsm['X_umap']

# %% [code]
adata.var_names

# %% [code]
results_file = './tabula_muris-scvr_prep.h5ad'
adata.write(results_file)

# %% [code]
x,y,z = adata.obsm['X_umap'][:,0], adata.obsm['X_umap'][:,1], adata.obsm['X_umap'][:,2]

x = np.array(x)
y = np.array(y)
z = np.array(z)

fig = plt.figure(figsize=(12,12), facecolor='w')
ax = fig.gca(projection='3d')
ax.scatter(x, y, z, s=0.1)

# %% [code]
pd.DataFrame(x).to_csv('3dplot.txt',sep='\n', index=False, header=False)

# %% [markdown]
Depending on the desired annotations and genes to be visualized by the user, the following command can be executed to create *singlecellVR*-compatible .JSON object. 

`scvr -f ./tabula_muris-scvr_prep.h5ad -t scanpy -a ANNOTATIONS [-g GENES] [-o OUTPUT]`
```
