## Supplementary IPython Notebooks for "*singlecellVR*: interactive visualization of single-cell data in virtual reality": Supplementary_Notebook_S3B.ipynb

```
# %% [markdown]
### Supplementary Notebook S1B
---
### STREAM SCoPE2 analysis performed on a monoctye differentiation dataset [(Specht et al., 2020)](https://doi.org/10.1101/665307)

The processed result of this datasest is shown in **Supplementary Figure 1B** of the manuscript.

**Preprint**: [Single-cell proteomic and transcriptomic
analysis of macrophage heterogeneity](https://www.biorxiv.org/content/10.1101/665307v4.full.pdf) 

**Raw Data**: [link](https://scope2.slavovlab.net/docs/data#processed-single-cell-protein-data) 

# %% [code]
import stream as st
st.__version__

# %% [code]
st.set_figure_params(dpi=100,style='white',figsize=[5.4,4.8],
                     rc={'image.cmap': 'viridis','lines.markersize':4})

# %% [markdown]
### Read in data

# %% [code]
adata=st.read(file_name='SCoPE2_GeneNames.h5ad',
              workdir='./result_stream_SCoPE2_GeneNames_0805')

# %% [code]
adata.X = adata.X.toarray()

# %% [code]
adata.X

# %% [code]
adata.var_names_make_unique()

# %% [code]
adata.var

# %% [markdown]
### Preprocess
##### It is preprocessed data and there is no need to do filter and normalization again.

# %% [code]
### filter
#st.filter_cells(adata,min_n_features= 100)
#st.filter_features(adata,min_n_cells = 5)

# %% [code]
### Normalize gene expression based on library size  
#st.normalize(adata,method='lib_size')

### Logarithmize gene expression 
#st.log_transform(adata)

# %% [code]
adata

# %% [markdown]
### Feature selection

# %% [code]
st.select_top_principal_components(adata,n_pc=15,first_pc=True)

# %% [markdown]
### Dimension Reduction

# %% [code]
st.dimension_reduction(adata,method='mlle',feature='top_pcs',n_components=3,n_neighbors=15,n_jobs=8)

# %% [code]
st.plot_dimension_reduction(adata,color=['celltype'],
                            n_components=3,show_graph=False,show_text=False, plotly=True)

# %% [code]
st.plot_dimension_reduction(adata,color=['celltype','batch_digest','batch_sort', 'batch_chromatography'],
                            n_components=3,show_graph=False,show_text=False, fig_ncol= 2)

# %% [markdown]
### Trajectory Inference

# %% [code]
st.seed_elastic_principal_graph(adata,n_clusters=10)

# %% [code]
st.plot_dimension_reduction(adata,color=['celltype','batch_digest','batch_sort','batch_chromatography'],
                            n_components=3,show_graph=True,show_text=False, fig_ncol= 2)
st.plot_branches(adata,show_text=True)

# %% [code]
st.elastic_principal_graph(adata,epg_alpha=0.2,epg_mu=0.05,epg_lambda=0.05)

# %% [code]
st.plot_dimension_reduction(adata,color=['celltype','batch_digest','batch_sort', 'batch_chromatography'],
                            n_components=3,show_graph=True,show_text=False, fig_ncol= 2)
st.plot_branches(adata,show_text=False)

# %% [code]
st.plot_dimension_reduction(adata,color=['celltype'],show_graph=True,plotly=True)

# %% [markdown]
### Trajectory Visulization

# %% [markdown]
##### flat tree

# %% [code]
st.plot_flat_tree(adata,color=['celltype','batch_digest','batch_sort', 'batch_chromatography','S1_pseudotime'],
                  dist_scale=0.5,show_graph=True,show_text=True)

# %% [markdown]
##### Stream plot at single cell 

# %% [code]
st.plot_stream_sc(adata,root='S1',color=['celltype','S1_pseudotime'],
                  dist_scale=0.3,show_graph=True,show_text=True)

# %% [markdown]
##### stream plot

# %% [code]
st.plot_stream(adata,root='S1',color=['celltype','batch_digest','batch_sort', 'batch_chromatography'])

# %% [code]


# %% [markdown]
### Marker genes detection

# %% [markdown]
##### 1) detect marker genes for each leaf branch 

# %% [code]
st.detect_leaf_markers(adata,cutoff_zscore=1.0,
                      cutoff_pvalue=0.01, root='S1',n_jobs=4)

# %% [code]
adata.uns['leaf_markers_all'].head()

# %% [code]
adata.uns['leaf_markers'][('S1','S0')].head()

# %% [code]
adata.uns['leaf_markers'][('S0','S2')].head()

# %% [code]
adata.uns['leaf_markers'][('S0','S3')].head()

# %% [code]
adata.var_names

# %% [code]
st.plot_stream(adata,root='S1',color=['VIM'])

# %% [markdown]
##### 2) detect transition genes for each branch

# %% [code]
st.detect_transition_markers(adata,cutoff_spearman=0.4,
                             cutoff_logfc=0.25, root='S1',n_jobs=4)

# %% [code]
adata.uns['transition_markers'][('S1','S0')].head()

# %% [code]
st.plot_stream_sc(adata,root='S1',color=['VIM'],
                  dist_scale=0.3,show_graph=True,show_text=True)

# %% [markdown]
##### 3) detect marker genes that are differentially expressed between pairs of branches

# %% [code]
st.detect_de_markers(adata,cutoff_zscore=1,
                     cutoff_logfc=0.25, root='S1',n_jobs=4)

# %% [code]
adata.uns['de_markers_greater'][(('S0', 'S2'), ('S0', 'S3'))].head()

# %% [code]
st.plot_de_markers(adata)

# %% [markdown]
##### 4) detect cell population-specific markers

# %% [code]
st.detect_markers(adata,ident='celltype',cutoff_zscore=1.0,cutoff_pvalue=0.01)

# %% [code]
adata.uns['markers_celltype']['sc_u'].head()

# %% [code]
st.plot_stream_sc(adata,root='S1',color=['ACACB'],
                  dist_scale=0.3,show_graph=True,show_text=True)

# %% [code]
adata.uns['markers_celltype']['sc_m0'].head()

# %% [code]
st.plot_stream_sc(adata,root='S1',color=['LMNA'],
                  dist_scale=0.3,show_graph=True,show_text=True)

# %% [markdown]
## save results

# %% [code]
st.write(adata,file_name='stream_result_var.pkl')

# %% [code]
adata.obs_keys()

# %% [code]
st.save_vr_report(adata,ann_list=['S1_pseudotime','celltype','branch_id_alias'],
                  gene_list=adata.var_names.to_list())

# %% [markdown]
Depending on the desired annotations and genes to be visualized by the user, the following command can be executed to create *singlecellVR*-compatible .JSON object. 

`scvr -f ./stream_result_var.pkl -t STREAM -a ANNOTATIONS [-g GENES] [-o OUTPUT]`
```
